# A MATLAB Toolbox for Modeling Genetic Circuits in Cell-Free Systems

**DOI:** 10.1101/2020.08.05.237990

**Authors:** Vipul Singhal, Zoltan A. Tuza, Zachary Z. Sun, Richard M. Murray

## Abstract

We introduce a MATLAB based simulation toolbox, called txtlsim, for an *E. coli* based Transcription-Translation (TX-TL) system. This toolbox accounts for several cell-free related phenomena, such as resource loading, consumption, and degradation, and in doing so, models the dynamics of TX-TL reactions for the entire duration of batch-mode experiments. We use a Bayesian parameter inference approach to characterize the reaction rate parameters associated with the core transcription, translation and mRNA degradation mechanics of the toolbox, allowing it to reproduce constitutive mRNA and protien expression trajectories. We demonstrate the use of this characterized toolbox in a circuit behavior prediction case study for an incoherent feed-forward loop.

## I. Background

Synthetic biology is often described as an endeavour to integrate engineering principles into the process of designing novel biological systems. One of its key goals is the engineering of genetic circuits to function in a predictable manner, so that these circuits may be used to control cellular behavior [1], [2]. The design-build-test cycle of these circuits *in vivo*, however, can be time consuming and expensive. In disciplines like electrical and aeronautical engineering, the design-build-test cycle is accelerated with the help of rapid prototyping environments like breadboards and wind tunnels, and associated dynamics simulation software such as PSpice [3], [4] and XFlow. Analogously, it should be possible to accelerate the design of novel biological systems in synthetic biology using appropriate rapid prototyping tools.

Recently, cell-free protein synthesis systems have been used as prototyping platforms for the design of genetic circuits [5], [6]. Cell lysate-based systems in particular are made of three components: a cytoplasmic extract, a buffer containing energy and raw material molecules, and a solution containing the DNA that encodes the circuit to be implemented. One example of an *Escherichia coli* (*E. coli*) based cell-free system is the TX-TL (transcription-translation) system [7], [8], [9].

Cell-free systems have numerous advantages that make them suitable as prototyping platforms in synthetic biology. First, since the DNA encoding the circuit is not constrained by the need for replication, restrictions due to plasmid selection markers and antibiotic compatibility are lifted. This allows for the rapid exploration of genetic circuit variants by trying different combinations of DNA species. Second, time-consuming cloning and transformation steps may be bypassed by using linearized DNA in the form of polymerase chain reaction (PCR) products or *de novo* synthesized fragments, which speeds up the DBT cycle.

Examples of software for simulating general biochemical reaction networks include the TABASCO [10], COPASI [11], Bioscrape [12], and MATLAB Simbiology^®^. TABASCO and COPASI are fast general purpose solvers that can incorporate stochastic simulations into circuit dynamics. Bioscrape is a Python-based simulator that leverages the speed of Cython to perform fast stochastic simulations with time delays, cell lineage tracking, and Bayesian parameter inference. Simbiology^®^ is a MATLAB toolbox that follows the Systems Biology Markup Language (SBML) in its structure, in that it allows for the specification of the standard SBML features such as models, compartments, reactions, species, parameters, rules and events in an object-oriented manner. It works well with MATLAB’s ordinary differential equation solvers, local and global optimization methods, plotting tools, and other functionalities.

Examples of modeling studies specific to cell-free systems include the one performed by Stögbauer et al. [13], who described a minimal rate equation model of the PURExpress reconstituted gene expression system, and performed a fit of their model parameters to the experimentally measured time courses of both the expressed protein and mRNA. They did not, however, explore fitting their model to an *E. coli* extract. Karzburn and Noireaux [14], on the other hand, did fit models to protein and mRNA data from the TX-TL *E. coli* extract, but restricted their analysis to the first 60 minutes of system behavior. More recently, Moore et al. [15] have performed extensive characterization of parts in non-model bacteria extracts, such as those made from *Bacillus Mageterium*. None of these studies, however, provide a software toolbox or attempted to use their models to predict and validate the behavior of whole circuits.

In this paper, we build on our initial work in [16] to describe a MATLAB based software toolbox called txtlsim for prototyping genetic circuits in TX-TL. This toolbox comes with a library of parts that can be combined in different combinations to build circuit models that are predictive of the behavior of circuits *in vitro*. It does this by accounting for the loading of finite enzymatic resources, which can introduce complex interactions between otherwise non-interacting elements of genetic circuits [17]. Furthermore, it models the consumption of resources like nulceotides and amino acids, and does so without needing to model elongation processes at the single base or codon resolution. Another feature of the toolbox is its simple user interface, which requires only a few lines of code and allows the user to specify a genetic circuit at the level of promoters and genes, abstracting away all lower level interactions. Finally, we demonstrate a Markov chain Monte Carlo (MCMC) based multi-stage Bayesian inference procedure for characterizing the toolbox’s parameters, and use the characterized models to predict and experimentally validate the behavior of an incoherent feed-forward loop under a variety of experimental conditions.

The rest of this paper is organized as follows. In Section II, we describe the implementation details of the toolbox, including the biochemical equations, and show sample code for creating models of genetic circuits. Section III describes inference of the ‘core’ toolbox parameters, which are parameters associated with transcription, translation and RNA and resource degradation. In Section IV, we demonstrate the predictive capabilities of the toolbox using an incoherent feed-forward loop as an example circuit. In Section V, we discuss the results and possible future work. Finally, in Section VI, we describe the experimental and computational methods used for data collection and parameter inference.

## II. Implementation

In this section, we build upon our initial work in [16] to describe a software toolbox called txtlsim for simulating the behavior of genetic circuits in TX-TL. This toolbox is useful for modeling TX-TL experiments for two principal reasons. Firstly, modeling the reactions associated with TX-TL requires explicit accounting of the interactions of DNA and RNA with enzymatic resource species such as RNA polymerases, ribosomes, ribonucleases (RNases), and transcription factors. This quickly increases the complexity of the chemical reaction network being built, due to effects such as resource loading and retroactivity [17]. This complexity is further compounded by the need to account for nucleotide and amino acid consumption and degradation. The txtlsim toolbox abstracts away the need for the specification of these low level mechanisms and interactions, allowing the user to specify genetic circuits at the level of genes. Indeed, txtlsim comes with a library of parts that can be combined in different combinations to build genetic circuit models. Secondly, it is able to predict the behavior of a genetic circuit, based only upon the characterization of its constituent parts, as discussed in Section IV.

The txtlsim toolbox is written using MATLAB Simbiology^®^, enabling its models to be compatible with MATLAB’s visualization, simulation, and parameter estimation capabilities. Genetic circuits are specified using a set of input strings that specify DNA, small molecules, and other miscellaneous species. The toolbox then generates a deterministic mass action kinetics model of this circuit’s mechanics in TX-TL. A typical TX-TL model, specified by the user at the resolution of whole DNA, mRNA and protein species, comprises mechanisms for transcription, translation, RNA degradation, transcriptional regulation and the eventual inactivation of TX-TL itself. Other mechanisms, such as linear DNA and protein degradation or gamS-mediated protection against nucleases, may also be included [18].

### A. User Interface

We highlight several features of txtlsim. First, the toolbox requires only a few lines of code to generate a relatively complex chemical reaction network model of TX-TL. For instance, the constitutive expression of green fluorescent protein (GFP) can be modeled using the code shown below.

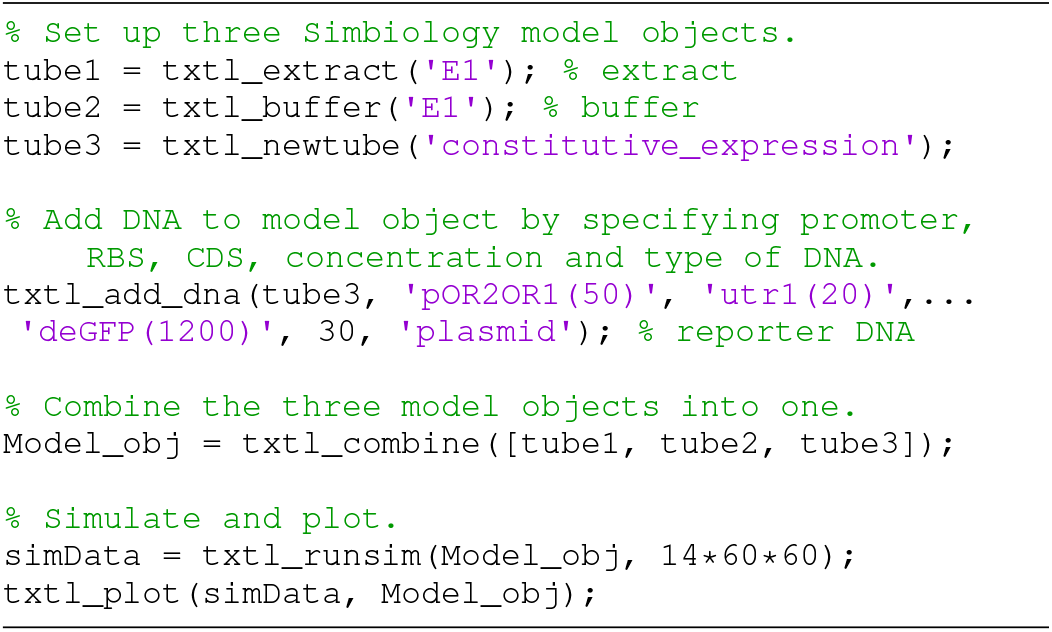

The set of commands shown in the snippet above mimic the actual experimental protocol of setting up a TX-TL experiment. The functions `txtl_extract` and `txtl_buffer` access extract and buffer specific parameter configuration files, selected by the input string `‘E1’` here, to set up two Simbiology^®^ model objects called `tube1` and `tube2`. The configuration files are user defined, and the parameters they contain can come from the literature, or from parameter inference performed on experimental data.

Next, the `txtl_newtube` and `txtl_add_dna` commands are used to initialize a new Simbiology^^®^^ model object and add DNA to this model object, respectively. In its most common use case, the `txtl_add_dna` function allows for specification of a promoter, an untranslated region and a coding sequence to form a transcriptional unit on the specified DNA, along with the concentration of the DNA added, and whether it is a linear fragment or plasmid DNA. For example, in the call to `txtl_add_dna` above, the promoter, ribosome binding site and coding sequence (CDS) are specified by the strings `‘pOR2OR1’, ‘utr1’` and `‘deGFP’`, respectively. These strings, each describing a component of the transcriptional unit, are used by `txtl_add_dna` to access a library containing code and parameter files associated with these components. These component files specify the reactions and species associated with each component, and encode interactions with other components. This allows txtlsim to automatically link different transcriptional units into a genetic circuit.

The `txtl_combine` command is used to combine the three model objects (`tube1, tube2` and `tube3`) into a model object, Mobj, which is then simulated using `txtl_runsim`, with the results plotted using `txtl_plot`. The flowchart in Figure 1 depicts the order these commands need to be specified in.

**Figure 1.**
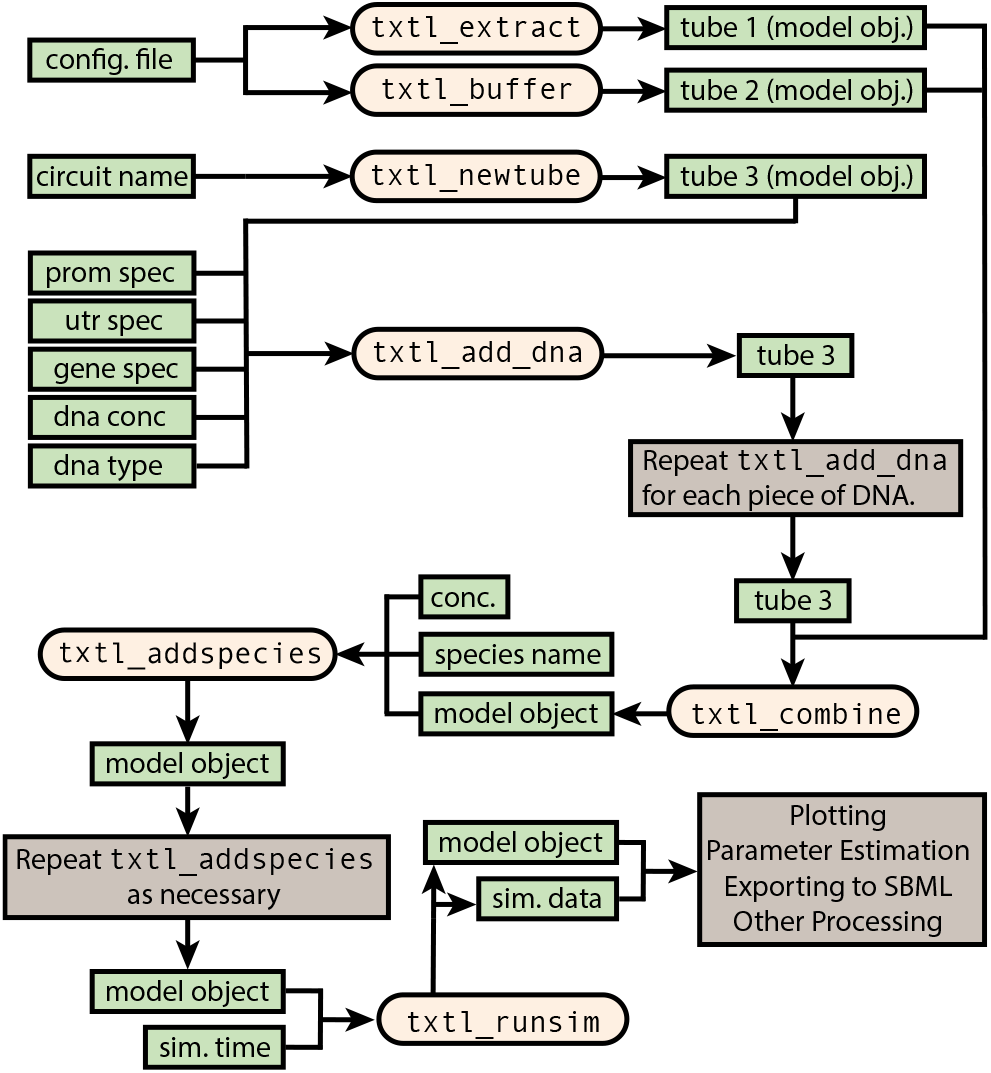
Flowchart of the user level code. The `txtl_add_dna` command is the main command that is used to specify the DNA to be added to the model. This allows for all the reactions and species associated with that DNA to be set up in the model. The model is contained in a Simbiology^®^ model class object, and is simulated using the `txtl_runsim` command.

Figure 2 shows the result of the `txtl_plot` command, which is arranged into three panels. The top panel shows the protein species that exist within the TX-TL system. Bottom left panel shows RNA (solid) and DNA (dashed) dynamics. The bottom right panel (normalized to 1) shows the dynamics of enzymatic and consumable resources. The enzymatic resources are ribosomes, RNA polymerases, and RNases; the consumable species in the model are the four nucleotides (ATP, GTP, CTP, and UTP), and amino acids (AA).

**Figure 2.**
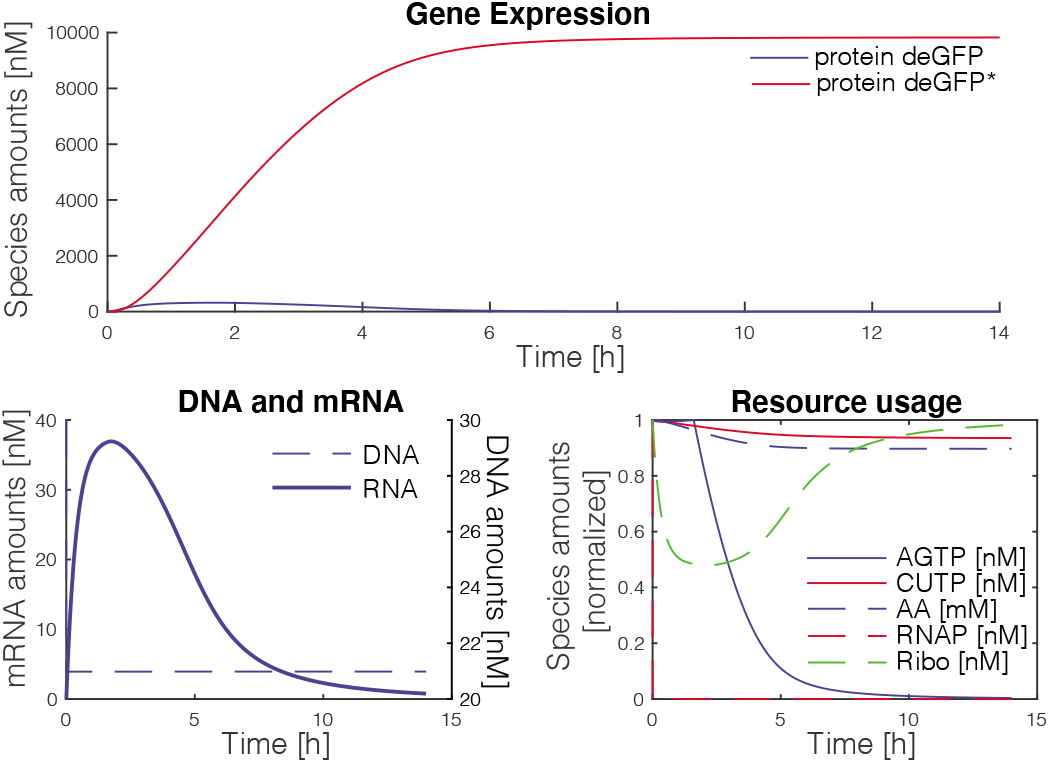
Standard output of txtlsim. Top: Gene expression profiles for unfolded (deGFP) and folded (deGFP*) reporter protein. Bottom left: left axis: mRNA profile, right axis: free DNA profile. Bottom right: Normalized resource loading and consumption. AGTP = ATP + GTP, CUTP = CTP + UTP.

The plots in Figure 2 were generated using parameters found by fitting the deGFP and mRNA curves to corresponding data, as described in Section III.

### B. The Modeling Framework of the txtlsim Toolbox

In this section, we describe the typical reaction network generated by txtlsim when a transcriptional unit is expressed. More complex networks made out of multiple transcriptional units interacting via transcription factor mediated regulation are simply iterations of this canonical network, but coupled via enzymatic and consumable resources, and the relevant regulatory interactions. The species in the software toolbox may be divided into five broad categories: DNA, mRNA, proteins, miscellaneous species such as inducers or nucleotides, and the biochemical complexes formed by combining these in defined ways.

The species follow a naming convention, which allows for the automated decision making involved in the reaction network generation. This naming convention is described in Supplementary Section S1.2.

The main processes associated with each transcriptional unit are transcription, translation and RNA degradation (Supplementary Figure S1). Other processes include DNA and protein degradation, transcription factor mediated activation and repression, and inducer action.

#### 1) Transcription

Transcription is modeled as a four step process (RNA polymerase binding, nucleotide binding, elongation and termination) using the reactions

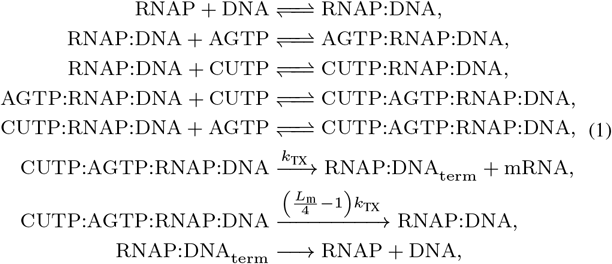

where the complex formed by two species, e.g. RNAP + DNA, is denoted as RNAP:DNA.

The catalytic machinery of transcription is lumped into a single species, denoted by RNAP. It is assumed to encompass RNA polymerases, sigma factors, and other cofactors. Transcription factors are modeled separately, because they are needed for transcriptional regulation, are user defined, and various distinct transcription factors may exist in a single circuit.

While all four nucleotides are used as raw materials for mRNA synthesis, GTP and ATP are also used as a source of energy for translation. Furthermore, they are modeled to undergo degradation about 1.5–3 hours after the initiation of the TX-TL experiment (as seen in [19] Figure 1B, where this degradation happens after 3 hours). Thus, the reactions CTP and UTP take part in are identical, but distinct from those that ATP and GTP are involved in, which are also identical. We have lumped these pairs of species into the so-called ‘CUTP’ and ‘AGTP’ species, respectively, where one molecule of CUTP models one molecule of CTP and one of UTP (similarly for AGTP).

After the binding of the catalytic and consumable species, the production of mRNA itself is divided into two reactions, an mRNA production reaction and a nucleotide consumption reaction. This latter reaction uses up AGTP and CUTP without producing mRNA, and is used to balance the consumption of nucleotides with the production of mRNA.

As an example, consider the transcription of an mRNA species that is a thousand base-pairs long. The toolbox assumes that each mRNA is composed of the four types of bases in equal proportions. Then, 250 molecules each of ATP, GTP, CTP and UTP are required for the transcription of this mRNA species. Thus, 250 units each of AGTP and CUTP are needed to transcribe the 1000 base-pair long mRNA molecule. Looking at the model in equation (1), we see that the mRNA production step consumes one unit each of AGTP and CUTP and produces one mRNA molecule. The consumption reaction also consumes one unit each of AGTP and CUTP, and does not produce an mRNA molecule. Thus, to consume 250 units each of AGTP and CUTP per mRNA produced (at quasisteady-state), we may set the rate of the consumption reaction to be *L*_m_/4 – 1 = 249 times the rate of the mRNA production step.

Finally, at the end of mRNA production, a termination complex RNAP:DNA_term_ forms, which then dissociates into RNAP and DNA in a separate reaction.

#### 2) Translation

Translation is modeled analogously, with the reactions

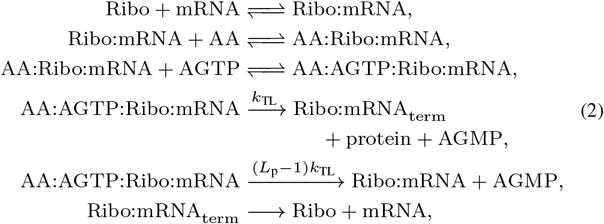

where *L*_p_ is the length of the protein in amino acids, and *k*_TL_ is the translation rate.

#### 3) RNA Degradation

RNA degradation is mediated by RNases, and is implemented as an enzymatic reaction,

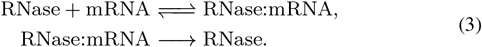

Similar binding and degradation reactions are set up for mRNA in its various bound forms, such as Ribo:mRNA, AA:Ribo:mRNA, etc., which result in the degradation of the mRNA and return of the remaining species to the species pool. The full set of these reactions is described in Supplementary Section S2.2.

#### 4) AGTP Regeneration System

The final core mechanism present in the toolbox models the AGTP regeneration system. This system has been shown to become inactivated after some time, leading to a degradation in its ability to express proteins [19], [20]. This can be seen in, for instance, Figure 2, where we observe that the inflection point for RNA production (bottom left panel) coincides with the inactivation of ATP regeneration system (bottom right panel).

We model this mechanism as a reversible degradationregeneration reaction involving AGTPand AGMP,

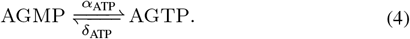

After *τ*_ATP_ seconds, the reverse (regeneration) reaction stops, leading to pure degradation of the energy resources in the system. The parameter *τ*_ATP_ must be estimated from experimental data, as described in Section III. In addition to ATP hydrolysis and depletion, [20], [21] discussed the depletion of other substrates (such as arganine, serine, cysteine and secondary energy sources like PEP and pyruvate) as having an effect on the inactivation of cell extracts. We simplify the modeling of these effects by lumping them with the AGTP degradation-regeneration mechanics.

#### 5) Other Reactions

Additionally, txtlsim can model linear DNA degradation mediated by RecBCD, which is a three subunit enzyme that unwinds DNA, and RecBCD sequestration by the GamS protein [18]. The TX-TL system has no innate protein degradation, and degradation of tagged proteins can be mediated by the ClpXP protease [22], [23] and transcription factor mediated regulation, among other mechanisms. For brevity, we describe just the transcriptional repression and induction reactions here. Repression by the dimerizable protein TetR and its sequestration by the inducer anhydrotetracycline (aTc) are modeled as

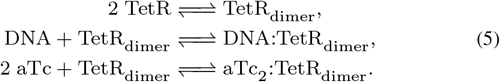

### C. Circuit Example

The incoherent feed-forward loop (IFFL) is a genetic circuit involving an activator, a repressor and a reporter (Figure 4A). Owing to the circuit’s network topology, the repression of the reporter is delayed with respect to its activation. The IFFL used in this paper uses LasR as an activator, expressed constitutively under the control of a pLac promoter. The repressor is TetR, which is under the control of an engineered pLas promoter. We also combined this las-activatable promoter with a tetO operator site to form the combinatorial promoter used to control the deGFP expression (Section VI-D). The code below shows the commands needed to set up the IFFL in txtlsim. This gene circuit will be used to demonstrate the predictive capabilities of the toolbox (Section IV and Supplementary Section S3).

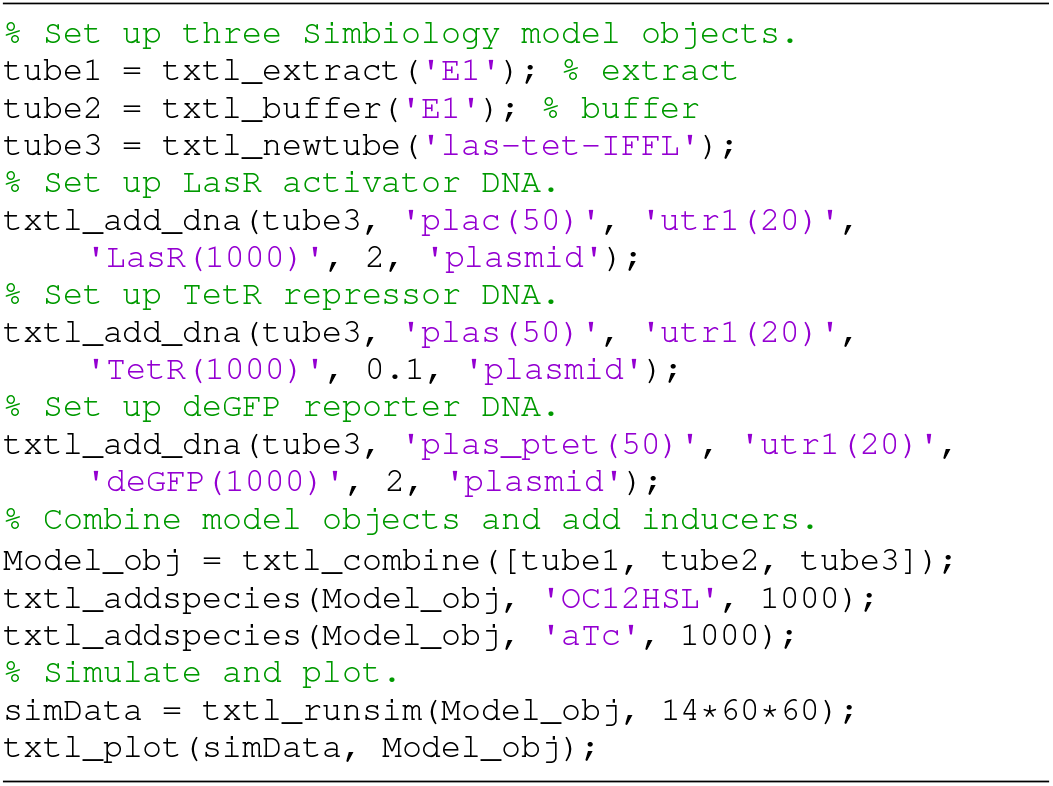

### D. Managing Chemical Reaction Network Complexity

In lower level specifications, such as Simbiology^®^, Bioscrape [12] or even directly stated ODEs, modeling the IFFL at the level of detail of the txtlsim toolbox would require several tens to over a hundred equations, all of which would need to be manually specified. Furthermore, modifying the network would entail manually updating the relevant equations, a process that is both time consuming and error prone. The txtlsim toolbox, on the other hand, allows the user to specify genetic circuits at a higher level of abstraction, allowing for rapid testing of different designs.

The rationale behind this number of equations to model even simple circuits is to account for the consumption of the limited pool of nucleotides and amino acids, and the loading of the finite catalytic and regulatory machinery (RNA polymerases, ribosomes, transcription factors, etc). The consumption and degradation of nucleotides and amino acids is thought to underlie the inactivation of the gene expression capability, and is therefore important to model to capture the full curves of TX-TL experiments. Coupling between different parts of a circuit, via the loading of enzymatic resources [17] or regulatory elements, has been shown to introduce unintended interactions between parts of genetic circuits in both TX-TL and *in vivo* [24]. The txtlsim toolbox incorporates these types of effects naturally, since it is built on mass action—as opposed to Michaelis-Menten or Hill—kinetics, allowing for enzymatic loading to be modeled by the explicit formation of complexes and a drop in the concentration of free enzyme. The use of such mass action kinetic models also means that the models are extensible, in the sense that once a species exists, new types of interaction can be added without modifying any of the existing equations—a property that does not hold for Michaelis-Menten or Hill kinetics in general. Finally, models created with txtlsim can be converted into SBML, and may be exported into any other SBML compatible environment for analysis.

In Supplementary Section S1.1, we describe the software architecture that allows for the automatic generation of these reactions and the interactions between them without the need for the user to specify them explicitly.

## III. Inferring the Parameters Associated with the Core Mechanisms in TX-TL

In this section, we estimate the parameters associated with the core mechanisms of TX-TL, such as transcription, translation, and RNA and resource degradation (equations (1)–(4)). Our parameter inference is performed in a Bayesian framework, with an MCMC sampler used to construct the posterior distribution of parameters, conditioned on the data and models (See Section VI, Materials and Methods). The experimental data used for estimating the core parameters is from [14], [25], and comprises fluorescence measurements of constitutive protein and mRNA expression, along with the degradation of spiked in mRNA (Figure 3, left column). More details about the data can be found in Supplementary Section S2.1.

**Figure 3.**
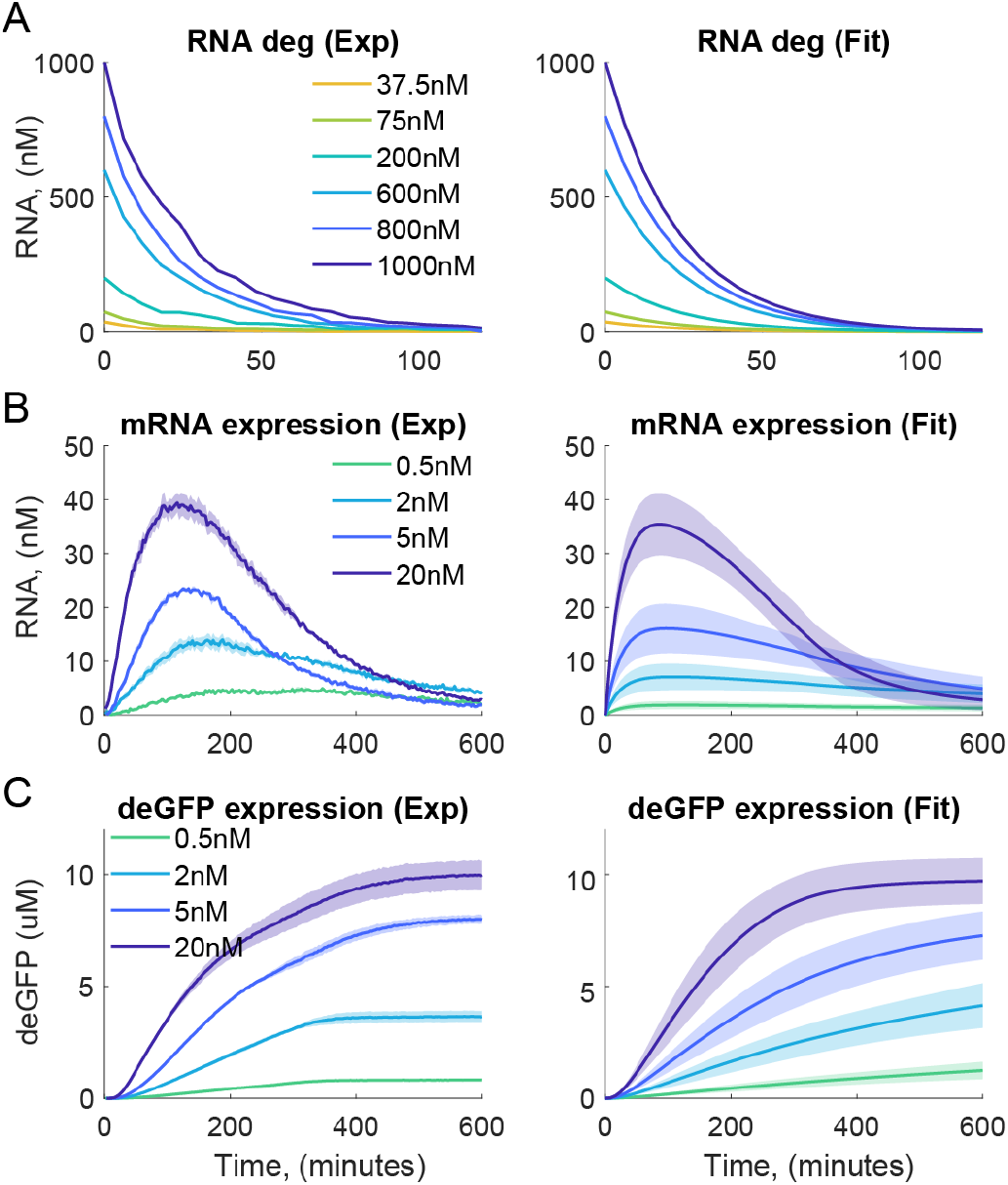
Estimating the core TX-TL parameters. Experimental data is from [14], [25]. Shaded regions indicate standard error over three replicates (left), and model simulations based on the inferred parameters (right). (A) Decay of purified deGFP-MGapt transcripts initiated at six different mRNA concentrations. (B) Transcription kinetics reported by a Malachite Green aptamer. (C) Translation kinetics reported by deGFP. Rows (B) and (C) show five different concentration of plasmid DNA that was added to each TX-TL master mix at the beginning of the experiment.

The column on the right of Figure 3 shows the results of fitting txtlsim models to this data. There are a total of 26 parameters in these models, of which several are non-identifiable [26], [27]. Unlike point estimation methods, MCMC based sampling gives an estimate of the entire joint parameter distribution, and can be used to gain insight into which parameters are non-identifiable and may therefore be fixed during parameter inference. Examples of such parameters include the forward reaction rates associated with reversible reactions, and in some cases, might even include the dissociation constants themselves. The forward reaction rates set the timescales at which fast reversible reactions reach quasi-steady-state, and were found to be non-identifiable once they were large enough for time-scale separation to be achieved. Some of the dissociation constants, especially those associated with reactions embedded deeper inside the reaction network, also had broad distributions, indicating possible non-identifiability. Examples of reactions with such dissociation constants include those involving nucleotides and amino acids binding to their respective transcription and translation complexes.

Table I and Supplementary Figures S2-S4 show five parameter inference runs involving models of constitutive expression and RNA degradation, and experimental data from [25]. In these runs, successively more parameters were fixed as part of an exploratory scheme designed to help elucidate the tradeoff between model fidelity and the computational tractability of the parameter inference procedure as a function of the number of free and fixed parameters. In Run 1, only the forward rates in the various reversible reactions were fixed, resulting in a 19-dimensional parameter space, which was too large to be searched efficiently. In Run 2, in addition to the forward reaction rates, the dissociation constants of the reactions involved in the binding of nucleotides and amino acids to the transcription and translation complexes were also fixed, leading to a more manageable search space. In the remaining runs, we successively increased the number of parameters that were fixed, allowing for the parameter space to be searched more efficiently. We found that Runs 3 and 4 were able to fit the data well while still allowing the parameter space to be searched relatively quickly.

**Table I.** Runs 1 - 5: Different combinations of parameters that were fixed during the core parameter inference. asterisks denote the parameters are fixed to the corresponding values at the nominal point.

| Parameter | nominal | Run 1 | Run 2 | Run 3 | Run 4 | Run 5 |
| --- | --- | --- | --- | --- | --- | --- |
| $TX_{cat}$ | 4.9 | est | est | est | est | est |
| $\tau_{atp}$ | 9.2 | est | est | est | * | * |
| $\delta_{atp}$ | -9.5 | est | est | est | * | * |
| $\alpha_{atp}$ | -3.9 | est | est | * | * | * |
| $pol_{Kd}$ | 9.5 | est | est | est | est | est |
| $pol_F$ | 1.5 | * | * | * | * | * |
| $pol_{term}$ | 3.3 | est | est | est | est | * |
| $n_{Kd1}$ | 2.9 | est | * | * | * | * |
| $n_{F1}$ | 0 | * | * | * | * | * |
| $n_{Kd2}$ | 14.0 | est | * | * | * | * |
| $n_{F2}$ | 0 | * | * | * | * | * |
| $RNAse_{Kd}$ | 9.2 | est | est | est | * | * |
| $RNAse_F$ | 0 | * | * | * | * | * |
| $RNAse_{cat}$ | -4.4 | est | est | est | est | * |
| $pol$ | 1.4 | est | est | est | est | est |
| $RNAse$ | 6.5 | est | est | est | est | * |
| $TL_{cat}$ | 0.5 | est | est | est | est | est |
| $GFP_{mat}$ | -6.1 | est | est | * | * | * |
| $Ribo_{Kd}$ | 11.2 | est | est | est | est | est |
| $Ribo_F$ | -0.2 | * | * | * | * | * |
| $aa_{Kd}$ | 6.6 | est | * | * | * | * |
| $aa_F$ | -0.3 | * | * | * | * | * |
| $TL_{n,Kd}$ | 14.5 | est | * | * | * | * |
| $TL_{n,F}$ | -1.2 | * | * | * | * | * |
| $Ribo_{term}$ | 5.4 | est | est | est | est | * |
| $Ribo$ | 7.3 | est | est | est | est | est |

The nominal parameter point shown in the first column provided the values to fix the parameters to, and was found using a combination of MCMC and manual tuning (details in Supplementary Section S2). Ultimately, the fits shown in Figure 3 were generated using the parameter fixing profile of Run 3, which had a small enough number of free variables to search over efficiently, while still being able to fit the models to the data. Supplementary Figure S5 shows the marginal distributions of the core parameters estimated during Run 3, using data from both [14], [25]. All parameter values are log-transformed (base-*e*).

## IV. Case Study: Experimentally Validated Prediction of Gene Circuit Behavior

The main role of the txtlsim toolbox is to be a high-fidelity simulator for the TX-TL system. This section highlights this role using a case study involving the prediction and experimental validation of the behavior of an IFFL.

First, we collected experimental data involving the components of an IFFL in five different experiments (Figure 4A, B and Table II, top half). Second, we estimated the parameters of the building blocks of IFFL, with some of the previously estimated core parameters (Supplementary Figure S5) providing the approximate ranges of values to search over, while others providing the values to fix the parameters to. Third, we assembled the whole gene circuit model of the IFFL using these characterized parts in txtlsim (Section II-C) and simulated it. We also collected experimental data about the dynamical behavior of the whole IFFL in TX-TL under five sets of experimental perturbations (Figure 5A, and lower part of Table II), and compared the simulated model with the experimental data (Figure 5B and C).

**Figure 4.**
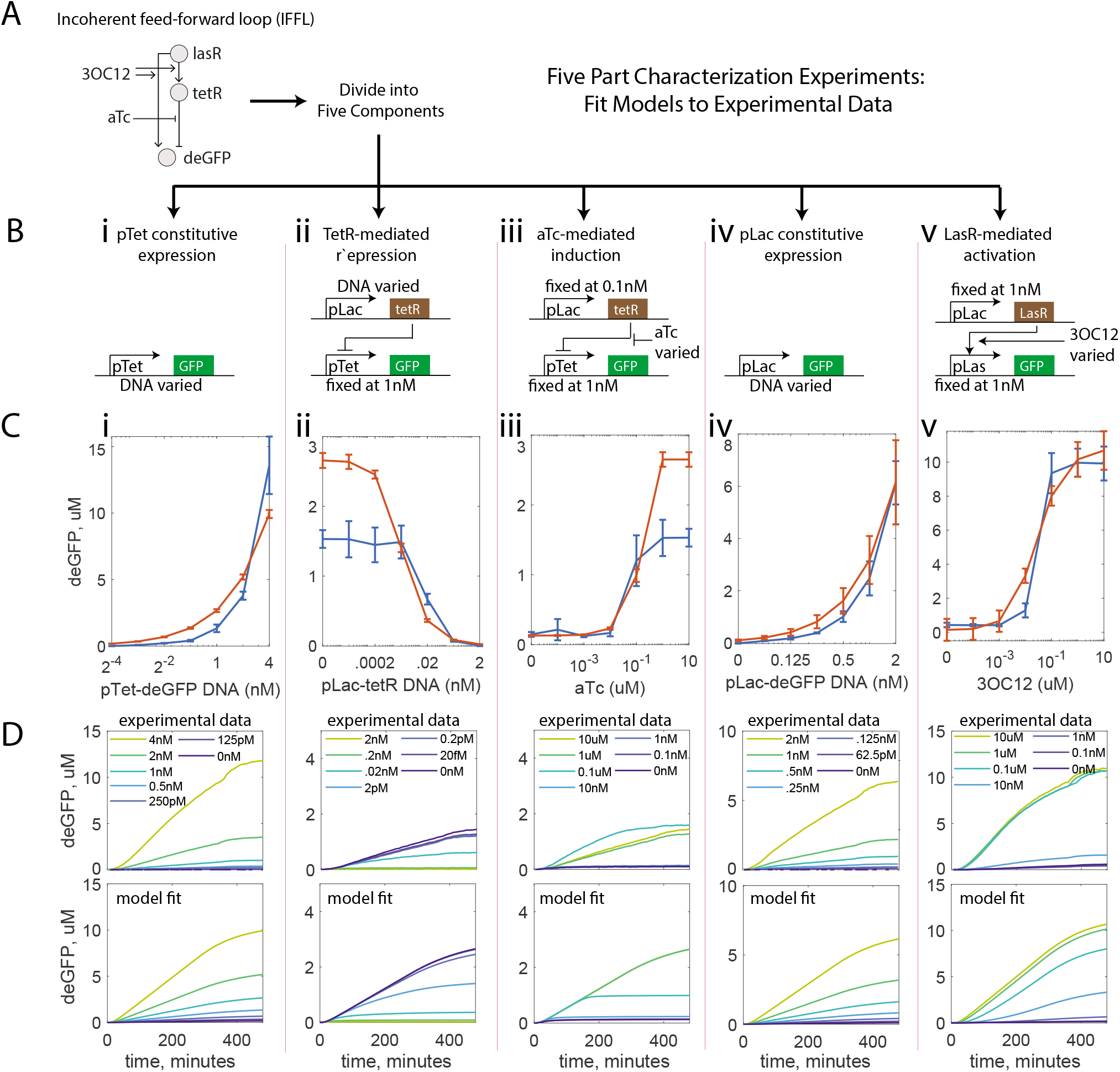
Part characterization and parameter fitting for the incoherent feed-forward loop (IFFL, schematic in A). (B) Schematics describing the five part characterization experiments used to infer the part-specific parameters. (C) Endpoint curves (mean, standard error at 480 min) corresponding to the experimental data (blue, n = 3) and corresponding parameter fitting trajectories (orange, n = 50, sampled from the posterior parameter distribution and simulated). The posterior distributions were generated by fitting the full time-course trajectories to the data (D).

**Table II.** IFFL part characterization and circuit behavior prediction experiments (Figures 4 and 5).

| Exp. Type | Experiment | Species Varied | Species Fixed |
| --- | --- | --- | --- |
| Characterization | Constitutive pTet expression | pTet-UTR1-deGFP: 4, 2, 1, 0.5, 0.25, 0.125 and 0.0625 nM |  |
|  | Constitutive pLac expression | pLac-UTR1-deGFP: 2, 1, 0.5, 0.25, 0.125, 0.0625 and 0.0313 nM |  |
| | TetR-mediated repression | pLac-UTR1-TetR: 2, 0.2, 0.02, 0.002, $2 \times 10^{-4}$, $2 \times 10^{-5}$ and 0 nM | pTet-UTR1-deGFP: 1 nM |
| | aTc-mediated induction | aTc: 10, 1, 0.1, 0.01, 0.001, $1 \times 10^{-4}$ and $1 \times 10^{-5}$ μM | pLac-UTR1-TetR: 0.1 nM<br>pTet-UTR1-deGFP: 1 nM |
| | 3OC12HSL-mediated induction | 3OC12 at 10, 1, 0.1, 0.01, 0.001, $1 \times 10^{-4}$ and $1 \times 10^{-5}$ μM | pLac-UTR1-LasR: 1 nM<br>pLas-UTR1-deGFP: 1 nM |
| Prediction | 3OC12 induction | 3OC12 at 10, 1, 0.1, 0.01, 0.001, $1 \times 10^{-4}$ and $1 \times 10^{-5}$ μM | Unless in the ‘Species Varied’ column:<br>pLac-UTR1-LasR: 1 nM<br>pLas-UTR1-TetR: 0.1 nM<br>pLas_tetO-UTR1-deGFP: 1 nM<br>aTc: 10 μM<br>3OC13: 1 μM |
| | LasR activation | DNA pLac-UTR1-LasR: 2, 1, 0.5, 0.25, 0.125, 0.0625 and 0.03125 μM | |
| | aTc induction | aTc: 10, 1, 0.1, 0.01, 0.001, $1 \times 10^{-4}$ and $1 \times 10^{-5}$ μM | |
| | TetR repression | DNA pLas-UTR1-TetR: 1, 0.1, 0.01, 0.001, $1 \times 10^{-4}$, $1 \times 10^{-5}$ and 0 μM | |
| | Reporter DNA | pLas_tetO-UTR1-deGFP: 4, 2, 1, 0.5, 0.25, 0.125 and 0.0625 μM | |

### A. Part Characterization

The part characterization experiments involved collecting data on the isolated behavior of the various components of the IFFL. The experiments we chose to characterize the components were pTet constitutive expression, TetR-mediated repression, aTc-mediated induction, pLac constitutive expression and 3OC12HSL-mediated induction (Figure 4B, and Table II, top half).

We used three models to fit the five IFFL characterization data sets shown in Table II. The constitutive pLac expression and the 3OC12HSL-mediated induction data sets had their own models, while the remaining three data sets: constitutive pTet expression, TetR-mediated repression and aTc-mediated induction only needed a single model, with three different sets of initial conditions accounting for their differences. Model equations and associated reaction rate parameters can be found in Supplementary Section S3.2.

The inference of the parameters associated with the parts of the IFFL was also performed in a consensus Bayesian inference framework (Section VI, Materials and Methods). In total, there were 42 parameters in the model, as shown in the first column of Table III. The first 26 parameters were those associated with the core mechanisms in the toolbox, and were described in Section III. The next 8 parameters were associated with the Tet-repression system, and the final 8 parameters were those associated with the Las-activation system.

**Table III.** Step-wise parameter inference strategy for IFFL part characterization.

| Stage | Init. | St. 1 | St. 2a | St. 2b | St. 2c | St. 2d | St. 2e | St. 2f |
| --- | --- | --- | --- | --- | --- | --- | --- | --- |
| Model | const. | const. | tet | tet<br>lac | tet<br>lac<br>las | tet<br>lac<br>las | tet<br>lac<br>las | tet<br>lac<br>las |
| $pol_{F, lac}$ (or p70) | 1.5 | * | * | * | * | * | * | * |
| $pol_{Kd, lac}$ (or p70) | 9.5 | 13.6 | * | free | free | free | free | free |
| $n_{F1}$ | 0 | * | * | * | * | * | * | * |
| $n_{Kd1}$ | 2.9 | * | * | * | * | * | * | * |
| $n_{F2}$ | 0 | * | * | * | * | * | * | * |
| $n_{Kd2}$ | 14.0 | * | * | * | * | * | * | * |
| $Ribo_F$ | -0.2 | * | * | * | * | * | * | * |
| $aa_F$ | -0.3 | * | * | * | * | * | * | * |
| $aa_{Kd}$ | 6.6 | * | * | * | * | * | * | * |
| $TL_{n,F}$ | -1.2 | * | * | * | * | * | * | * |
| $TL_{n,Kd}$ | 14.5 | * | * | * | * | * | * | * |
| $RNase_F$ | 0 | * | * | * | * | * | * | * |
| $GFP_{mat}$ | -6.1 | * | * | * | * | * | * | * |
| $\alpha_{atp}$ | -3.9 | * | * | * | * | * | * | * |
| $TX_{cat}$ | 4.9 | 2.4 | * | 2.3 | * | free | 3.1 | * |
| $pol_{term}$ | 3.3 | 4.4 | * | * | * | free | free | free |
| $pol$ | 1.4 | 1.6 | * | free | free | free | free | free |
| $TL_{cat}$ | 0.5 | 3.3 | * | 3.7 | * | free | 3.4 | * |
| $Ribo_{Kd}$ | 11.2 | 0.05 | * | * | * | * | * | * |
| $Ribo_{term}$ | 5.4 | 2.8 | * | * | * | free | free | free |
| $Ribo$ | 7.3 | 4.2 | * | free | free | free | free | free |
| $RNase_{Kd}$ | 9.2 | 15.6 | * | * | * | * | * | * |
| $RNase_{cat}$ | -4.4 | -0.2 | * | * | * | * | * | * |
| $RNase$ | 6.5 | 8.6 | * | 9.2 | * | * | * | * |
| $\tau_{atp}$ | 9.2 | 8.9 | * | 10.1 | * | 9.7 | * | * |
| $\delta_{atp}$ | -9.5 | -9.7 | * | * | * | * | * | * |
| $pol_{F, tet}$ | | | Fixed: 1.5 | * | * | * | * | * |
| $pol_{Kd, tet}$ | | | free | free | free | free | free | free |
| $rep_{Kd}$ | | | free | -2.7 | * | -0.5 | * | * |
| $rep_F$ | | | 1.3 | * | * | * | * | * |
| $aTc_{Kd}$ | | | free | -6.0 | * | free | -2.0 | * |
| $aTc_F$ | | | 1.6 | * | * | * | * | * |
| $dim_{Kd}$ | | | -10.0 | * | * | * | * | * |
| $dim_F$ | | | 1.4 | * | * | * | * | * |
| $pol_{F, las}$ | | | | | fixed: 0 | * | * | * |
| $pol_{Kd, las}$ | | | | | free | free | free | free |
| $3OC12_{Kd}$ | | | | | free | 13.0 | * | * |
| $3OC12_F$ | | | | | fixed: 0 | * | * | * |
| $pLas-pol_{TF, F}$ | | | | | fixed: 0 | * | * | * |
| $pLas-pol_{TF, Kd}$ | | | | | free | free | free | free |
| $pLas_{TF, F}$ | | | | | fixed: 0 | * | * | * |
| $pLas_{TF, Kd}$ | | | | | free | free | free | free |

Table III summarizes the multi-stage parameter estimation scheme that we employed to search the parameter space. This approach was needed because the 42 dimensional space was too large to be searched efficiently. All columns except the one labeled ‘Init’ describe a parameter inference run in terms of which parameters were estimated, and which ones were fixed. There are four types of symbols in this table: Asterisks, the phrase ‘fixed: p’ (where p is a numerical parameter value), numerical values, and the word ‘free’. Asterisks indicate that the value was the same as the value in the previous column. The phrase fixed: 1.5 means that that parameter value was fixed to 1.5 at that stage. A numerical value indicates that that parameter was freely estimated at that stage, and that value was picked from the resulting distribution (jointly with any other such fixed values) and fixed in the next stage (therefore, every numerical value is necessarily followed by an asterisk). Finally, the word ‘free’ means that that parameter was freely estimated at that stage, but no value was picked for fixing in the next stage (and is therefore never followed by an asterisk in the next column). As in the previous section, all parameter values are log-transformed (base-*e*).

The first 26 (core) parameters are specified by the ‘Init.’ (initialization) and ‘St. 1’ (Stage 1) columns of the table. Initialization refers to the initial parameter point found using MCMC and manual parameter tuning in Section III. The numbers shown in the Stage 1 column correspond to a particular parameter point from the distribution found in Run 3 during the core inference stage.

In Stage 2a, the constitutive pTet expression, TetR-mediated repression and aTc-mediated induction circuits described in Table II and Figure 4 were characterized. In addition to the core parameters, there were 8 additional Tet-system associated parameters in the model. Stage 2b had the same models, but with some of the core and Tet-system parameters re-estimated. In stage 2c, the 3OC12HSL-mediated induction (activation of the pLas promoter by the LasR activator) system was introduced, along with 8 new parameters associated with this model. The forward rates are fixed to a value of zero (in log_*e*_-space), and the parameters marked ‘free’ are estimated. In stages 2d-f, we estimated different combinations of parameters, and in doing so, explored the trade-off between model fidelity and the computational tractability of the MCMC runs. The characterization results from the probability distribution resulting from Stage 2f are shown in Figure 4, and the pairwise marginal probability distributions are shown in Supplementary Figure S7. The fitting trajectories and parameter distributions resulting from Stage 2d are shown in Supplementary Figure S8.

### B. Model Prediction and Experimental Validation

Using the part-specific parameters inferred in the previous section, we created *in silico* predictions of the behavior of the whole IFFL, and compared these to corresponding TX-TL data. We measured the behavior of the IFFL under five sets of perturbations away from a nominal IFFL. This nominal condition was: pLac-UTR1-LasR at 1nM; IPTG at 1mM (sequestering any native LacI in the extract); the LasR inducer 3OC12HSL at 1 μM; the repressor DNA pLas-UTR1-TetR at 0.1 nM; the reporter DNA plas_tetO-UTR1-deGFP at 1nM; and the TetR inducer aTc at 10 μM.

With this nominal IFFL, we collected the deGFP expression levels under perturbations of 3OC12HSL, the LasR DNA, aTc, the TetR DNA and the deGFP DNA, as listed in lower half of Table II. The results of these experiments are shown in Figure 5B and C. The blue curves in Figure 5B show the expression levels of the deGFP under the various perturbations at 480 minutes (i.e., end-point measurements). The error bars are the standard errors of three technical replicates. Figure 5C, top row, shows the trajectories of the full 480 minutes of these experiments.

**Figure 5.**
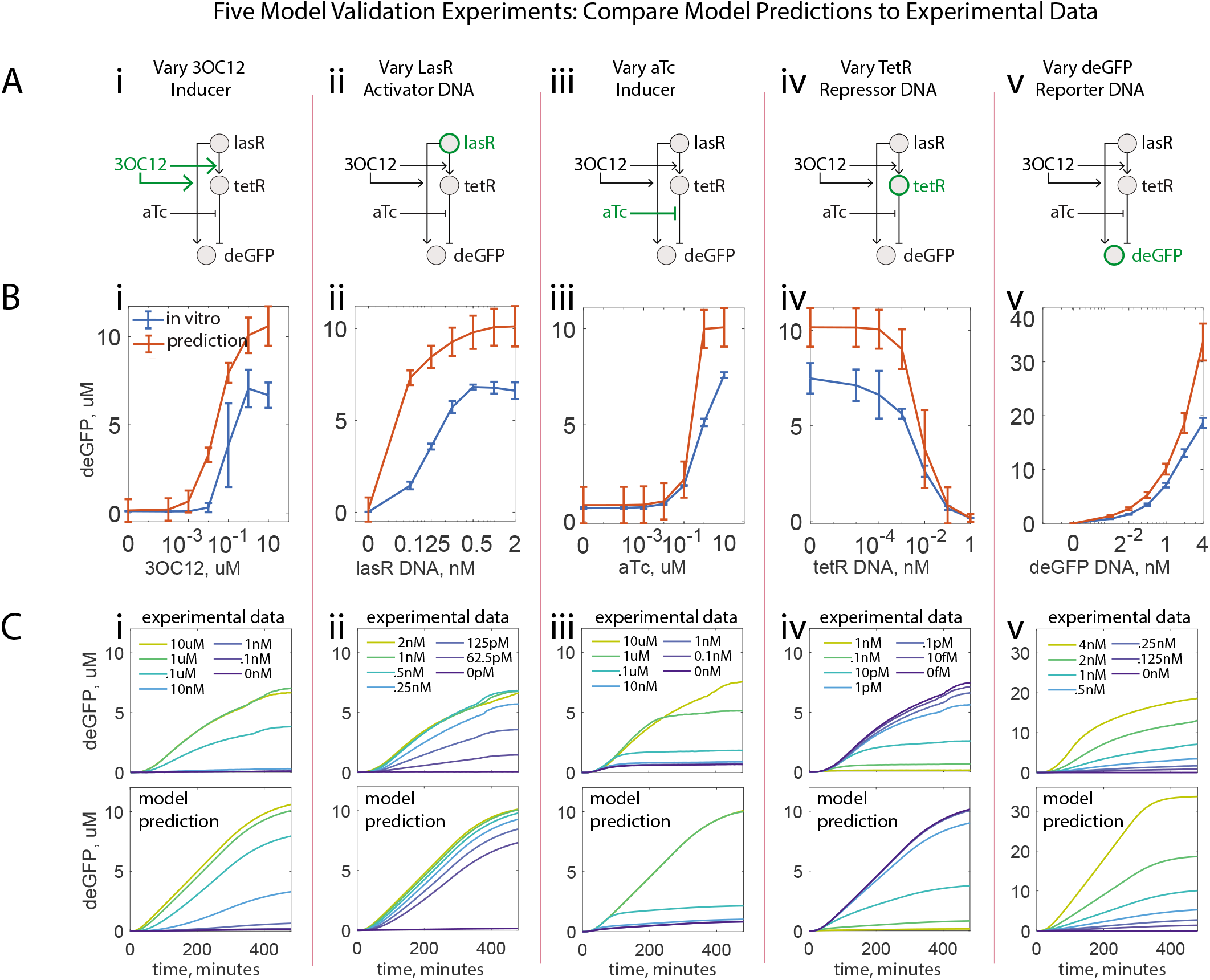
Model predictions and experimental validation for the incoherent feed-forward loop (IFFL). (A) Schematics describing the five perturbations of the IFFL that were used for the validation of model predictions. (B) IFFL behavior under these perturbations. The nominal IFFL conditions are described in the main text. Endpoint measurements of mean and standard error for experimental (blue, n = 3) and predicted (orange, n = 50) values (t = 480 min). (E) Corresponding experimental and model prediction trajectories.

The model prediction trajectories were generated by sampling points from the joint posterior parameter distribution resulting from Stage 2f in Table III, and simulating the IFFL model at each of these parameter points. The orange curves in Figure 5B are the mean and standard errors of the deGFP species concentration at 480 minutes for 50 of these trajectories. Similarly, the predictions shown in the bottom row of Figure 5C are the mean trajectories of the same 50 trajectories.

## V. Discussion

Synthetic biology is an attempt at incorporating engineering principles into the design of novel biological functions. These principles include the standardization of parts, the principled composition of these parts into larger systems, the use of abstraction layers to decouple phenomena at different scales, and the use of rapid prototyping and predictive modeling to speed up the engineering process.

The rapid prototyping paradigm has been implemented in synthetic biology using cell-free systems like TX-TL, which have become a tool for testing genetic circuits *in vitro*. In this paper, we have described an *in silico* modeling toolbox called txtlsim to accompany TX-TL. This toolbox is built using MATLAB Simbiology^^®^^ and closely mimics the species, reactions and chemical reaction network dynamics of TX-TL.

In particular, txtlsim has a number of specific features that make it suited as a tool for circuit behavior prediction in TX-TL. Firstly, it explicitly models the usage of enzymes like RNA polymerases and ribosomes using mass action kinetics. Unlike Hill or Michaelis-Menten kinetics, this accounts for the loading of enzymatic or consumable resources once the binding reactions involving these resource species are specified. Accounting for the usage of amino acids and nucleotides usually requires modeling transcription and translation at the single base or amino acid resolution [28], [29], making the genetic circuit models large and unwieldy. We avoid this by modeling these processes in a single step reaction, while accounting for the resource usage by a separate consumption reaction, with a reaction rate that maintains the correct stoichiometry.

This toolbox also has a library of parts that can be composed to build a spectrum of circuits. As discussed in this study, the circuit models built out of part models that have characterized reaction rate parameters should be predictive of the *in vitro* behavior of the corresponding circuits. We demonstrated this using the IFFL circuit. We first characterized the parameters associated with the core transcription, translation and mRNA degradation mechanics of the toolbox, followed by the parameters associated with the parts of the IFFL. The characterized models were then combined into a model of the IFFL, and we verified that the predicted model behavior matched the corresponding experimental data.

Parameter inference of the individual circuit parts may be performed using MATLAB’s in-built optimization tools, or using the MCMC based consensus Bayesian inference tools provided with this toolbox. The Bayesian approach gave estimates of the joint distribution of the part-parameters, conditioned on the data, models, and any fixed parameters. Visualizing the (marginalized) probability densities can be used to indicate which parameters might be non-identifiable, or even co-varying with other parameters. This allowed us to fix the values of highly non-identifiable parameters, making otherwise computationally intractable inference problems tractable. Furthermore, the ability to visualize the co-variation [30] between parameters allows us to perform this fixing while respecting the relationships between parameters. These considerations were crucial for performing the inference of both the core model parameters and the circuit-part specific parameters.

Despite the features present in this version of txtlsim, there are several directions it can be extended in. Most simply, various new circuit features may be added to the library of parts, such as antisense RNA mediated transcriptional regulation [6] or integrase mediated DNA recombination [31]. Capabilities for accounting and correcting for extract batch variation [30], [32], studying the identifiability of the circuits, or for predicting the *in vivo* behavior from the *in vitro* behavior may also be added, although these tasks present significant research challenges.

As the field of synthetic biology matures, we expect computational modeling to play in increasingly predictive role in the design of genetic circuits, just as it has played in electrical, mechanical and aeronautical engineering.

## VI. Materials and Methods

### A. TX-TL Extract and Buffer Preparation

Preparation and execution of TX-TL was according to previously described protocols [9], with a modification of the strain used to ExpressIQ (New England Biolabs).

Briefly, the cells were grown to an OD_600_ of 1.5, pelleted and washed. They were then lysed using bead beating, and centrifuged to remove the beads and cell debris. The supernatant was incubated at 37° C for 80 min, and then centrifuged to remove endogenous nucleic acids. The supernatant was dialyzed against a pH8.2 buffer containing Mg-glutamate, K-glutamate, Tris, and DTT. Finally, the extract was centrifuged and the supernatant was flash-frozen in liquid nitrogen and stored at −80°C.

The buffer had the following components: 9.9 mg/mL protein, 9.5 mM Mg-glutamate, 95 mM K-glutamate, 0.33 mM DTT, 1.5 mM each amino acid except leucine, 1.25 mM leucine, 50 mM HEPES, 1.5 mM ATP and GTP, 0.9 mM CTP and UTP, 0.2 mg/mL tRNA, 0.26 mM CoA, 0.33 mM NAD, 0.75 mM cAMP, 0.068 mM folinic acid, 1 mM spermidine, 30 mM 3-PGA, 2% PEG-8000.

Both the extract and buffer were stored at −80°C in separate tubes, with enough volume for seven reactions per tube.

### B. TX-TL Experiment

A 384-well microplate (Nunc) was used for the experiments, and the appropriate concentrations and volumes of DNA and inducers to be used in each reaction were calculated using the spreadsheets provided in [9]. The extract and buffer were thawed for 20 min on ice, mixed in the prescribed ratios, and pipetted into each well being used in the microplate, which was also placed on ice. The DNA was then added to each well according to the spreadsheet. All the pipetting was done to avoid bubbles, the plate was sealed, and spun at 4000g for 45 s at 4°C to distribute the mix evenly at the bottom of the wells and remove any bubbles that might have been introduced. The plate was placed in a Synergy H1/MF microplate reader (Biotek). Settings used for deGFP measurement were: excitation/emission 485 nm/515 nm, at gain 61, measured every 8 min for 8 hours.

### C. Plasmid construction

DNA was cloned using standard molecular biology procedures [33], [34], [35] and propagated in a JM109 recA-lacIQ (Zymo Research) strain for purification. Small scale purifications were done by miniprep (PureYield, Promega) followed by a PCR purification for desalting (QiaQuick, Qiagen). Large scale purifications were done by midiprep or maxiprep (NucleoBond Xtra Midi or NucleoBond Xtra Maxi, Macherey-Nagel). All plasmids were isolated in stationary phase and sequenced before use.

### D. IFFL Part Screening in TX-TL

We first characterized a LasR-responsive promoter, pRsaL (Porig), from previously published work by testing its ability to express deGFP in the presence of LasR and 3OC12HSL (Supplementary Figure S6A) [36]. While the promoter was responsive to LasR, the *V*_max_ of the promoter was lower than anticipated and the dynamic range was under 6-fold (Supplementary Figure S6B, C). To find a more robust part, we used TX-TL to screen four more promoters taken from the Registry of Standard Biological Parts or from RNAseq data of known responsive elements [37]. Out of our screen, P1 showed a 7-fold improved *V*_max_ over Porig and a 29fold dynamic range (Supplementary Figure S6B, C). We also characterized the basal leakiness of the promoters without LasR present (Supplementary Figure S6D). Finally, we confirmed the result from our extract was generalizable by testing all 5 promoters for *V*_max_ in 11 independently made extracts using the same method [9] but over four *E. coli* strains (Supplementary Figure S6E). We then used P1 for the downstream LasR-responsive promoter due to its high *V*_max_. To engineer a TetR-repressible, LasR activatable combinatorial promoter, we tried two placements of the tetO operator sites (Supplementary Figure S6F-I) and characterized the response of these variants under aTc activation.

### E. Modeling Framework

We assume mass action kinetics, along with a well stirred, constant temperature and volume assumption on our reactions. This allows us to model the chemical equations as a set of ordinary differential equations (ODEs) with the reaction rate parameters and the unknown initial concentrations as the parameters of the system. Formally, we define an experiment 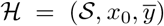 to be the execution of a system 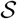 under initial conditions x_0_ and output measurements 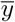, where the bar denotes the assumption that experimental data reflects the ground truth. With each experiment, we associate an initialized, parametrized ODE model *M_i_*, with the general structure

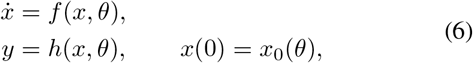

where the state vectors, which encode the species concentrations, are 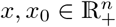, and are assumed to exist for all *t* ≥ 0. The parameter vector symbol is 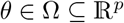, where Ω is the set of all possible parameter points of interest. The output is denoted 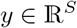, where S is the number of output variables.

### F. Experiment Ensemble and Consensus Parameter Inference

In general, we have multiple experiments informing some common set of parameters, where a given experiment may not inform every parameter, but every parameter is informed by at least one experiment.

Consider an ensemble of experiments 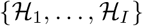, and an associated ensemble of models {*M*_1_,…, *M_I_*}, where model *M_i_* with parameters 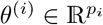 captures the evolution of the system in experiment 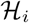 under the specified initial conditions.

The different experiments range over different doses (initial conditions), replicates, and genetic circuits (systems). The data are collected at a given sampling rate, which discretizes the trajectories. For the *i*th experiment, the discretization of 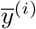 may be written as a matrix, 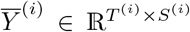, where we note that the number of time points *T*^(*i*)^ and measured output variables *S*^(*i*)^ will in general depend on *i*. We concatenate the set of these matrices into a block

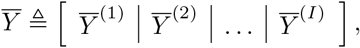

matrix with the appropriate padding of zeros when the number of time points differ between experiments.

We collect all the parameters from the ensemble into a *master vector*, 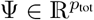, counting a parameter that appears in multiple *θ*^(*i*)^’s only once, so that 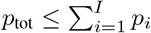. The individual parameter vectors can be related to the master vector via a binary *membership matrix* Γ ∈ {0, 1}^*p*_tot_×*I*^, where the (*k, j*) - th entry is 1 if the *k*th element of Ψ is present in *θ*^(*j*)^, and 0 otherwise.

If we group the individual parameter vectors into a matrix 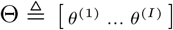, and consider diag(Ψ) as the square matrix with the elements of Ψ on the diagonal, and zeros elsewhere, then the matrix equation Θ = diag(Ψ) · Γ relates the master vector to the individual models’ parameters via the membership matrix. Consensus parameter inference is described visually in Figure 6.

**Figure 6.**
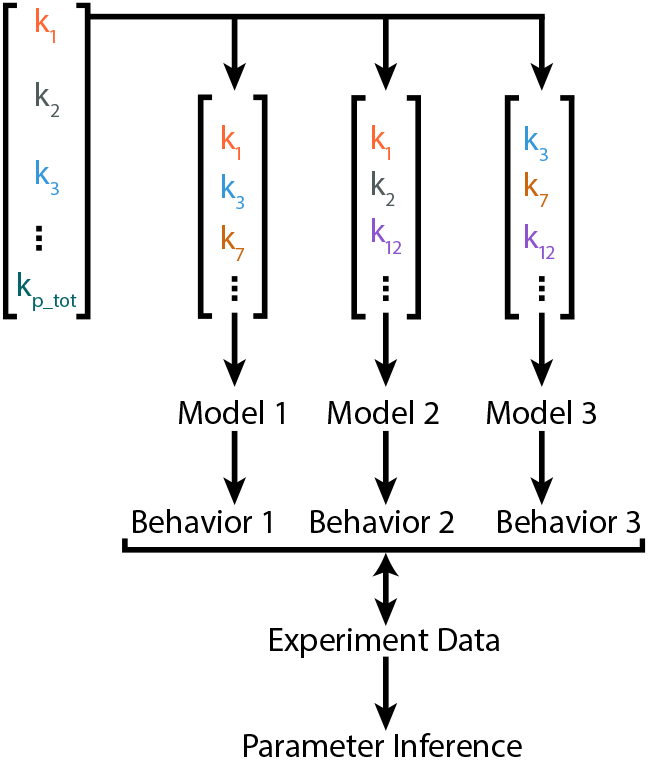
Parameter overlap in the consensus parameter inference problem.

### G. Bayesian Parameter Inference

We wish to determine the probability density of the parameters, conditioned on the experiments, models, and consensus pattern,

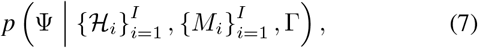

where the data for experiment *i* are included by virtue of being an element of the experiment tuple, 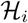. In what follows, we simplify notation by replacing 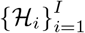 with the data matrix 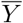, as defined in the previous section. We also drop Γ and 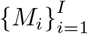, though these are assumed. Then, Bayes rule gives

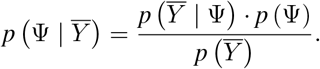

We assume the prior to be uninformative (uniform within a hypercube, and zero outside). The likelihood function involves data from all the experiments, measured species, replicates, and time points, and is defined as

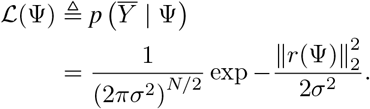

The vector *r*(Ψ) is defined as

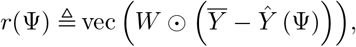

with the ⊙ symbol denoting the elementwise (Hadamard) product, and vec denoting the column-wise reshaping of a matrix into a vector. *N* denotes the total number of data points in the output of the models. The model predictions 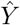 depend on the parameter values Ψ, and are arranged the same way as 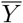. The matrix *W* has the same shape as 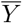, and contains weights to normalize for the different magnitudes of different output variables. For example, the concentrations of proteins are often much greater than those of mRNA, and fitting performance can be improved greatly by normalizing these data sets using *W*. Finally, we follow the standard practice of working with log-probabilities, which improves both the speed and stability of the numerical computations ([38], chapter 22).

In general, there is no analytical description of the parameter distributions associated with biochemical reaction networks. The standard approach is to use Markov chain Monte Carlo (MCMC) methods to sample from the desired parameter distribution, and construct an approximation to this distribution. This is done by constructing a Markov chain that has this distribution as its stationary distribution. It does this by performing a random walk in parameter space, such that the probability of being in a given region is proportional to the desired probability density in that region [39]. We used the MATLAB implementation from [40] of the emcee sampler [41], [42], with some modifications for better walker initialization and handling of numerical ill-conditioning.

## Code Availability

The code for this toolbox is available at https://github.com/vipulsinghal02/txtlsim_buildacell, along with multiple tutorials, and scripts to generate the figures in the paper and the supplementary information.

## Author’s contributions

RMM conceived of the txtlsim toolbox, and VS, ZAT and RMM wrote it. ZZS conceived of the experiments and collected the data, and VS conceived of and performed the parameter inference strategy. VS wrote the manuscript. All authors edited it.

## Acknowledgements

We thank Kyle Martin, Shaobin Guo and Clarmyra Hayes for laboratory assistance, and Vincent Noireaux and Michael Jewett for advice on extract preparation methods. We thank Anandh Swaminathan, William Poole, and Samuel Clamons for useful discussions about the computational methodology.

This material is based upon work supported in part by the Defense Advanced Research Projects Agency (DARPA/MTO) Living Foundries program, contract number HR0011-12-C-0065 (DARPA/CMO); the Air Force Office of Scientific Research, grant number FA9550-14-1-0060; and DARPA SBIR contract W911NF-16-P-0003 (Caltech subcontract from Synvitrobio, Inc.).

VS was supported by the National Science Scholarship (NSS-PhD), Agency for Science, Technology, and Research (A*STAR), Singapore. ZAT was supported by the Fulbright Scholarship. ZZS was supported by a UCLA/Caltech Medical Scientist Training Program fellowship, and a National Defense Science and Engineering Graduate fellowship.

The views and conclusions contained in this document are those of the authors and should not be interpreted as representing officially policies, either expressly or implied, of the Defense Advanced Research Projects Agency or the U.S. Government.

## References

[1] T.S. Gardner, C. R. Cantor, and J. J. Collins, “Construction of a genetic toggle switch in Escherichia coli,” Nature, vol. 403, no. 6767, pp. 339–342, Jan. 2000.

[2] M. B. Elowitz and S. Leibler, “A synthetic oscillatory network of transcriptional regulators,” Nature, vol. 403, no. 6767, pp. 335–338, Jan. 2000.

[3] L. W. Nagel and D. Pederson, “Spice (simulation program with integrated circuit emphasis),” EECS Department, University of California, Berkeley, Tech. Rep. UCB/ERL M382, Apr 1973.

[4] “Xflow: Fluids simulations to improve real-world performance,” Software, Dessault Systems, 2020.

[5] H. Niederholtmeyer, Z. Z. Sun, Y. Hori, E. Yeung, A. Verpoorte, R. M. Murray, and S. J. Maerkl, “Rapid cell-free forward engineering of novel genetic ring oscillators,” eLife, vol. 4, p. e09771, Oct. 2015.

[6] M. K. Takahashi, J. Chappell, C. A. Hayes, Z. Z. Sun, J. Kim, V. Singhal, K. J. Spring, S. Al-Khabouri, C. P. Fall, V. Noireaux, R. M. Murray, and J. B. Lucks, “Rapidly Characterizing the Fast Dynamics of RNA Genetic Circuitry with Cell-Free Transcription–Translation (TX-TL) Systems,” ACS Synthetic Biology, Mar. 2014.

[7] J. Shin and V. Noireaux, “Efficient cell-free expression with the endogenous E. Coli RNA polymerase and sigma factor 70,” Journal of Biological Engineering, vol. 4, p. 8, Jun. 2010.

[8] J. Shin and V. Noireaux, “An E. coli Cell-Free Expression Toolbox: Application to Synthetic Gene Circuits and Artificial Cells,” ACS Synthetic Biology, vol. 1, no. 1, pp. 29–41, Jan. 2012.

[9] Z. Z. Sun, C. A. Hayes, J. Shin, F. Caschera, R. M. Murray, and V. Noireaux, “Protocols for Implementing an Escherichia coli Based TX-TL Cell-Free Expression System for Synthetic Biology,” Journal of Visualized Experiments: JoVE, no. 79, Sep. 2013.

[10] S. Kosuri, J. R. Kelly, and D. Endy, “TABASCO: A single molecule, base-pair resolved gene expression simulator,” BMC Bioinformatics, vol. 8, no. 1, p. 480, Dec. 2007.

[11] S. Hoops, S. Sahle, R. Gauges, C. Lee, J. Pahle, N. Simus, M. Singhal, L. Xu, P. Mendes, and U. Kummer, “COPASI—a COmplex PAthway Simulator,” Bioinformatics, vol. 22, no. 24, pp. 3067–3074, Dec. 2006.

[12] A. Swaminathan, V. Hsiao, and R. M. Murray, “Quantitative Modeling of Integrase Dynamics Using a Novel Python Toolbox for Parameter Inference in Synthetic Biology,” bioRxiv, p. 121–152, Mar. 2017.

[13] T. Stögbauer, L. Windhager, R. Zimmer, and J. O. Rädler, “Experiment and mathematical modeling of gene expression dynamics in a cell-free system,” Integrative Biology, vol. 4, no. 5, pp. 494–501, May 2012.

[14] E. Karzbrun, J. Shin, R. H. Bar-Ziv, and V. Noireaux, “Coarse-Grained Dynamics of Protein Synthesis in a Cell-Free System,” Physical Review Letters, vol. 106, no. 4, p. 048104, Jan. 2011.

[15] S. J. Moore, J. T. MacDonald, S. Wienecke, A. Ishwarbhai, A. Tsipa, R. Aw, N. Kylilis, D. J. Bell, D. W. McClymont, K. Jensen, K. M. Polizzi, R. Biedendieck, and P. S. Freemont, “Rapid acquisition and model-based analysis of cell-free transcription–translation reactions from nonmodel bacteria,” Proceedings of the National Academy of Sciences, vol. 115, no. 19, pp. E4340–E4349, May 2018, iSBN: 9781715806118 Publisher: National Academy of Sciences Section: PNAS Plus.

[16] Z. A. Tuza, V. Singhal, J. Kim, and R. M. Murray, “An in silico modeling toolbox for rapid prototyping of circuits in a biomolecular “breadboard” system,” in 52nd IEEE Conference on Decision and Control, Dec. 2013, pp. 1404–1410.

[17] A. Gyorgy and D. Del Vecchio, “Limitations and trade-offs in gene expression due to competition for shared cellular resources,” in 53rd IEEE Conference on Decision and Control. IEEE, 2014, pp. 5431–5436.

[18] Z. Z. Sun, E. Yeung, C. A. Hayes, V. Noireaux, and R. M. Murray, “Linear DNA for Rapid Prototyping of Synthetic Biological Circuits in an Escherichia coli Based TX-TL Cell-Free System,” ACS Synthetic Biology, vol. 3, no. 6, pp. 387–397, Jun. 2014.

[19] V. Noireaux, R. Bar-Ziv, and A. Libchaber, “Principles of cell-free genetic circuit assembly,” Proceedings of the National Academy of Sciences, vol. 100, no. 22, pp. 12 672–12 677, 2003.

[20] D.-M. Kim and J. R. Swartz, “Prolonging cell-free protein synthesis with a novel ATP regeneration system,” Biotechnology and Bioengineering, vol. 66, no. 3, pp. 180–188, 1999.

[21] M. C. Jewett and J. R. Swartz, “Substrate replenishment extends protein synthesis with an in vitro translation system designed to mimic the cytoplasm,” Biotechnology and Bioengineering, vol. 87, no. 4, pp. 465–472, Aug. 2004.

[22] T. A. Baker and R. T. Sauer, “ClpXP, an ATP-powered unfolding and protein-degradation machine,” Biochimica Et Biophysica Acta, vol. 1823, no. 1, pp. 15–28, Jan. 2012.

[23] Z. Z. Sun, J. Kim, V. Singhal, and R. M. Murray, “Protein degradation in a TX-TL cell-free expression system using ClpXP protease,” bioRxiv, p. 019695, May 2015, publisher: Cold Spring Harbor Laboratory Section: New Results.

[24] D. Del Vecchio, A. J. Ninfa, and E. D. Sontag, “Modular cell biology: retroactivity and insulation,” Molecular Systems Biology, vol. 4, p. 161, Feb. 2008.

[25] Dan Siegal-Gaskins, Z. A. Tuza, J. Kim, V. Noireaux, and R. M. Murray, “Gene Circuit Performance Characterization and Resource Usage in a Cell-Free “Breadboard”,” ACS Synthetic Biology, vol. 3, no. 6, pp. 416–425, Jun. 2014.

[26] O.-T. Chis, J. R. Banga, and E. Balsa-Canto, “Structural Identifiability of Systems Biology Models: A Critical Comparison of Methods,” PLoS ONE, vol. 6, no. 11, p. e27755, Nov. 2011.

[27] A. Raue, C. Kreutz, T. Maiwald, J. Bachmann, M. Schilling, U. Klingmüller, and J. Timmer, “Structural and practical identifiability analysis of partially observed dynamical models by exploiting the profile likelihood,” Bioinformatics, vol. 25, no. 15, pp. 1923–1929, Aug. 2009.

[28] S. Arnold, M. Siemann, K. Scharnweber, M. Werner, S. Baumann, and M. Reuss, “Kinetic modeling and simulation of in vitro transcription by phage T7 RNA polymerase,” Biotechnology and Bioengineering, vol. 72, no. 5, pp. 548–561, Mar. 2001.

[29] R. Algar, T. Ellis, and G.-B. Stan, “Modelling the burden caused by gene expression: an in silico investigation into the interactions between synthetic gene circuits and their chassis cell,” arXiv:1309.7798 [q-bio], Sep. 2013, arXiv: 1309.7798.

[30] V. Singhal and R. M. Murray, “Transforming Data Across Environments Despite Structural Non-Identifiability,” in 2019 American Control Conference (ACC), Jul. 2019, pp. 5639–5646.

[31] V. Hsiao, Y. Hori, P. W. Rothemund, and R. M. Murray, “A populationbased temporal logic gate for timing and recording chemical events,” Molecular Systems Biology, vol. 12, no. 5, p. 869, May 2016.

[32] A. Gyorgy and R. M. Murray, “Quantifying resource competition and its effects in the tx-tl system,” in 2016 IEEE 55th Conference on Decision and Control (CDC), 2016, pp. 3363–3368.

[33] D. G. Gibson, L. Young, R.-Y. Chuang, J. C. Venter, C. A. Hutchison Iii, and H. O. Smith, “Enzymatic assembly of DNA molecules up to several hundred kilobases,” Nature Methods, vol. 6, no. 5, pp. 343–345, May 2009.

[34] A. Sarrion-Perdigones, E. E. Falconi, S. I. Zandalinas, P. Juárez, A. Fernández-del Carmen, A. Granell, and D. Orzaez, “GoldenBraid: An Iterative Cloning System for Standardized Assembly of Reusable Genetic Modules,” PLOS ONE, vol. 6, no. 7, p. e21622, Jul. 2011.

[35] C. Engler, R. Kandzia, and S. Marillonnet, “A One Pot, One Step, Precision Cloning Method with High Throughput Capability,” PLOS ONE, vol. 3, no. 11, p. e3647, Nov. 2008.

[36] A. Tamsir, J. J. Tabor, and C. A. Voigt, “Robust multicellular computing using genetically encoded NOR gates and chemical ‘wires’,” Nature, vol. 469, no. 7329, pp. 212–215, Jan. 2011.

[37] O. Wurtzel, D. R. Yoder-Himes, K. Han, A. A. Dandekar, S. Edelheit, E. P. Greenberg, R. Sorek, and S. Lory, “The Single-Nucleotide Resolution Transcriptome of Pseudomonas aeruginosa Grown in Body Temperature,” PLOS Pathogens, vol. 8, no. 9, p. e1002945, Sep. 2012.

[38] D. J. C. MacKay, Information Theory, Inference & Learning Algorithms. New York, NY, USA: Cambridge University Press, 2002.

[39] N. Christensen, R. Meyer, and A. Libson, “A Metropolis–Hastings routine for estimating parameters from compact binary inspiral events with laser interferometric gravitational radiation data,” Classical and Quantum Gravity, vol. 21, no. 1, p. 317, Jan. 2004.

[40] A. Grinsted, “Gwmcmc: An implementation of the goodman and weare mcmc sampler for matlab,” GitHub Repository, March 2015.

[41] J. Goodman and J. Weare, “Ensemble samplers with affine invariance,” Communications in Applied Mathematics and Computational Science, vol. 5, no. 1, pp. 65–80, Jan. 2010.

[42] D. Foreman-Mackey, D. W. Hogg, D. Lang, and J. Goodman, “emcee: The MCMC Hammer,” Publications of the Astronomical Society of the Pacific, vol. 125, no. 925, pp. 306–312, Mar. 2013, arXiv: 1202.3665.

