## Supplemental Information for "A MATLAB Toolbox for Modeling Genetic Circuits in Cell-Free Systems"

#### Contents

**S1 Architectural Details of the Toolbox** 2
S1.1 Automated Reaction Network Generation 2
S1.2 Species Naming Convention 3
S1.3 Directory Structure 4

**S2 Inferring the Core TX-TL parameters** 4
S2.1 Experimental Data 5
S2.2 Model Equations 5
S2.3 Parameter Inference 7

**S3 Inferring Incoherent Feed-forward Loop Part Parameters** 8
S3.1 Experimental Data 8
S3.2 Model Equations 8
S3.3 Parameter Inference 14
S3.4 Parameter Consensus Pattern 14

### S1 Architectural Details of the Toolbox

#### S1.1 Automated Reaction Network Generation

In Section II, we gave an overview of how a user may set up a simple circuit using a few lines of code in `txtlsim`. In this section, we go into further details of how the software sets up the model. We start with a walk-through of what each command does, along with a discussion of some of the architectural features of the toolbox. Figure 1 shows the flow of the code the user specifies to set up a model.

The commands `txtl_extract` and `txtl_buffer` are used to initialize the extract specific parameters and species. These functions set up a `txtl_reaction_config` class object that contains methods and properties to manage most of the core parameters in the model. The properties of the `txtl_reaction_config` class object are set by a configuration file stored in the `config` directory.

The command `txtl_add_dna` is the workhorse of the network generation phase of the toolbox, and is discussed in some detail here. It takes a model object as its first input, followed by a promoter specification string `promsepc`, a UTR specification string `utrspec`, a CDS specification `cdsspec`, a numerical DNA concentration, and a DNA type specification string as inputs (Table S1). Generically, a call to this function takes the form,

```
txtl_add_dna(model_object, promspec, utrspec,cdsspec, DNAconc, DNAtype),
```

where the `promsepc`, `rbssepc` and `cdsspec` strings are used to access component files of the same names in the component directory. These files contain all the relevant information pertaining to the promoter, RBS or CDS being specified, including the reactions it is involved in and the associated parameters.

**Table S1** Inputs to the `txtl_add_dna` command. The parenthetical arguments within the specifications are optional, and if they are not specified, then default values from the component configuration files are used. The DNA concentration can be any nonnegative numerical value, and the DNA type must be either `'linear'` or `'plasmid'`.

| Input | Syntax | Example |
|---|---|---|
| `model_object` | Simbiology® model object | `tube3` |
| `promspec` | string(optional numeric) | `'pOR2OR1(50)'` |
| `utrspec` | string(optional numeric) | `'UTR1(40)'` |
| `cdsspec` | string(optional numeric) | `'TetR(650)'` |
| `DNAconc` | numeric | 20 |
| `DNAtype` | string | `'linear'` |

The `txtl_add_dna` command performs the following actions: call the component function files for the promoter, the UTR and the CDS, followed by a function to set up mRNA degradation species and reactions, followed by DNA and protein degradation, if present. The promoter file sets up reactions and species (depending on the mode) associated with TF mediated regulation and transcription. Similarly, the UTR function file sets up ribosome binding reactions and other reactions associated with translation. The model per transcriptional unit (promoter, UTR and CDS) is summarized in Figure S1.

Returning to the user level code, once all the `txtl_add_dna` commands have been specified, the model objects specified by `tube1`, `tube2` and `tube3` are combined by the `txtl_combine` command, wich simply adds the species and reactions from the three model objects into a single model object, and scales the concentrations of the species to model the resulting change in volume. The resulting model object, `Model_obj`, can be

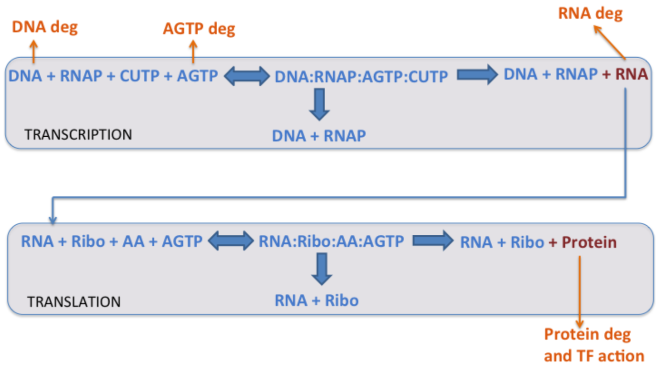


**Figure S1** A summary of the main mechanics that each transcriptional unit (TU) can be involved in.

simulated by `txtl_runsim`. Note that even if simulation is not the immediate goal, one call to `txtl_runsim` must be performed, since this is where the reactions in the model are set up. After the call to `txtl_runsim`, we have a fully defined model object, and a `simData` class object (a class in the Simbiology toolbox) containing the results of the simulation. These objecs may be used for further simulations, parameter inference, and visualization of the species trajectories, or be exported to other platforms via SBML.

#### S1.2 Species Naming Convention

In this section, we describe the species naming convention used by the toolbox, which parses these name strings when making decisions about how to generate the reaction network. Table S2 gives an overview of the naming conventions used for proteins, RNA and DNA.

**Table S2** Species naming conventions

| Species Type | Convention | Example |
|---|---|---|
| DNA | `DNA <promspec>--<utrspec>--<cdsspec>` | ‘DNA thio-junk-pTet--utr1--TetR’<br>‘DNA pTet--utr1--TetR-lva’ |
| RNA | `RNA <utrspec>--<cdsspec>` | ‘RNA utr1--TetR’<br>‘RNA att1-utr1--TetR-lva’ |
| protein | `protein <cdsspec>` | ‘protein TetR’<br>‘protein TetR-lva’ |

Here, `promspec`, `utrspec` and `cdsspec` are the promoter, untranslated region (UTR) and coding sequence specifications respectively. The specifications are separated by the double hyphen ‘--’, and within each specification, we may have various types of modifiers, separated by the single hyphen. Examples of these modifiers include `junk` DNA on the promoter to protect against DNA degradation, or `lva` protein degradation tags on the coding sequence. The miscellaneous species include inducers like anhydrotetracycline (aTc) or Isopropyl beta-D-1-thiogalactopyranoside (IPTG), core species like ribosomes (`ribo`), RNA polymerases (`RNAP`), `RecBCD` and `RNase` nucleases, etc., and resources like amino acids (`AA`) and grouped nucleotide species (`AGTP`, `CUTP`). Larger complexes may be formed using these elementary species by using the colon to join different species. For example, the complex comprising the 3OC12 inducer, the LasR protein, the DNA containing the pLas promoter and the RNA polymerase is given by `RNAP:DNA pLas_ptet--utr1--deGFP:OC12HSL:protein LasR`.

##### S1.3 Directory Structure

The main directory of the toolbox is called `trunk`. The key subdirectories in this directory are shown in Table S3. The core directory contains most of the source code of the network generation part of the toolbox. It contains user end functions like `txtl_add_dna` and hidden functions such as `txtl_mrna_degradation` or `txtl_transcription`. The `config` directory contains comma separated value (.csv) configuration files containing parameters associated with extracts and buffers. Each extract and buffer has its own configuration file, which is populated using calibration data collected for that extract and buffer. The `components` directory acts as a library of parts (such as promoters, UTRs and CDSs) from which genetic circuits may be constructed. It contains code (.m) and parameter configuration (.csv) files for these parts. Promoters in this library can be of an activatable, repressible or combinatorial nature. Some promoters, like the arabinose induced pBAD promoter may be repressed by a transcription factor (AraC in this case) when it is not bound to its inducer, and activated by the transcription factor when it is bound to the inducer. UTRs, such as the ribosome binding site, have their own files. Finally, CDSs can include reporters, repressors, activators, sigma factors, kinases, phosphatases or proteases, to name a few. All three classes of components can be extended in a straightforward manner to include new components.

**Table S3** Directory structure of the Toolbox

| **Directory** | **Description** |
|---|---|
| core | Core functions of the toolbox, such as `txtl_add_dna` or `txtl_transcription` |
| `config` | Extract and buffer configuration files (.csv). These contain parameters like transcriptional elongation rate, or the initial concentration of RNA polymerases or nucleotides corresponding to a given extract. |
| `components` | Component (promoter, UTR and CDS) files. This directory contains both code (.m) and parameter configuration (.csv) files. |
| `mcmc_simbio` | MCMC toolbox for Simbiology®. This toolbox allows for Bayesian parameter inference to be performed on the parameters of Simbiology® models with consensus over many model-data set pairings. |
| `examples` | Examples for the modeling toolbox. Includes examples from constitutive gene expression, to the incoherent feedforward loop and the genetic toggle. |
| `doc` | Contains the user manual and associated files. |

#### S2 Inferring the Core TX-TL parameters

We used experimental data from the literature to estimate biochemical reaction rate parameters associated with transcription, translation, RNA degradation and nucleotide degradation and regeneration. We denote these the 'core' mechanisms in the toolbox, and describe the inference of the parameters associated with them in this section.

The estimation of the core parameters had two goals. First, we wished to obtain an approximate characterization of the mechanics of the toolbox that allowed users to start using the toolbox immediately. Users can, for example, use the characterized toolbox to explore resource loading trade-offs under different experimental conditions. Second, we wanted to obtain approximate distributions of the values of the core parameters so that the non-identifiable parameters could either be fixed in the subsequent whole gene circuit characterization stage, or we could define appropriate bounds of the parameter space to be searched.

#### S2.1 Experimental Data

The experimental data were a combination of data from References [2] and [5]. Figures 1, S1 and S2 of Reference [2] provide trajectories of the constitutive expression of reporter protein (deGFP) and fluorescent RNA (Malachite Green bound RNA aptamer), and that of the degradation of spiked in fluorescent RNA. With 20 nM of constitutively expressed DNA, the maximum RNA levels observed were approximately 400 nM, and the deGFP levels were approximately 18 μM. The RNA degradation half life was found to be approximately 16–18 min. This data differed somewhat from that in Reference [5] (possibly due to batch effects, [4, 8], where, with 30 nM of plasmid DNA, the observed mRNA steady state was 20–30 nM 40–80 min after initiation of the reaction, and about 10 μM of reporter protein at the end of the reaction.

We chose to scale the trajectories from Reference [2] to make the data similar to that in Reference [5], and use the resulting 'composite' dataset for the core parameter inference. The mRNA and protein trajectories from Reference [2] were scaled down by a factor of 10 and 1.8 respectively. We verified that the original unmodified data could also be fit to the models. Indeed, as described in Section S2.3 below, we first fit the model to the unmodified data from [2] (Supplementary Figures S2-S4, and Runs 1 - 5 in Table I), and then picked one of the runs (Run 3) from this set as the basis for fitting the model to the composite data (Figure 3 and Supplementary Figure S5).

#### S2.2 Model Equations

The biochemical equations in the core model can be divided into those associated with transcription, translation, RNA degradation and miscellaneous reactions.

We associate a forward rate and a dissociation constant with each reversible reaction, with the understanding that the reverse reaction rate is given by the product of these two parameters. In this formulation, the forward rate can be thought to be setting the *timescale* of the reaction, and the dissociation constant the relative *strength* of the (un)binding reaction. In general, we found that most of the forward rate parameters could be fixed to values close to one (0 in log space), and the results of the simulations were insensitive to their exact values. Other parameters that were fixed, such as the dissociation constants for the binding of nucleotides to the DNA-RNAP complexes ($n_{Kd1}$, $n_{Kd2}$), the binding of amino acids or AGTP to the RNA-ribosome complex ($aa_{Kd}$, $TL_{n,Kd}$), the deGFP maturation rate or the AGTP regeneration rate were also fixed, because they were highly non-identifiable, and we found that the behavior of the system was insensitive to their exact values, or that other parameters could compensate for fixing of these parameters.

*Transcription reactions*

$$\text{RNAP} + \text{DNA} \underset{\text{pol}_{\text{Kd}}}{\overset{\text{pol}_{\text{F}}}{\rightleftharpoons}} \text{RNAP:DNA}, \qquad \text{Pol. Binding,}$$

$$\text{RNAP:DNA} + \text{AGTP} \underset{\text{n}_{\text{Kd1}}}{\overset{\text{n}_{\text{F1}}}{\rightleftharpoons}} \text{AGTP:RNAP:DNA}, \qquad \text{ATP-GTP binding,}$$

$$\text{RNAP:DNA} + \text{CUTP} \underset{\text{n}_{\text{Kd1}}}{\overset{\text{n}_{\text{F1}}}{\rightleftharpoons}} \text{CUTP:RNAP:DNA}, \qquad \text{CTP-UTP Binding,}$$

$$\text{CUTP:RNAP:DNA} + \text{AGTP} \underset{\text{n}_{\text{Kd2}}}{\overset{\text{n}_{\text{F2}}}{\rightleftharpoons}} \text{CUTP:AGTP:RNAP:DNA}, \qquad \text{ATP-GTP binding,}$$

$$\text{AGTP:RNAP:DNA} + \text{CUTP} \underset{\text{n}_{\text{Kd2}}}{\overset{\text{n}_{\text{F2}}}{\rightleftharpoons}} \text{CUTP:AGTP:RNAP:DNA}, \qquad \text{CTP-UTP binding,}$$

$$\text{CUTP:AGTP:RNAP:DNA} \xrightarrow{k_{\text{TX}}} \text{RNAP:DNA}_{\text{term}} + \text{mRNA}, \qquad \text{TX Elongation,}$$

$$\text{CUTP:AGTP:RNAP:DNA} \xrightarrow{\left(\frac{Lm}{4}-1\right)k_{\text{TX}}} \text{RNAP:DNA}, \qquad \text{Consumption,}$$

$$\text{RNAP:DNA}_{\text{term}} \xrightarrow{\text{pol}_{\text{term}}} \text{RNAP} + \text{DNA}, \qquad \text{Pol. Release.}$$

*Translation reactions*

$$\text{Ribo} + \text{mRNA} \underset{\text{ribo}_{\text{Kd}}}{\overset{\text{ribo}_{\text{F}}}{\rightleftharpoons}} \text{Ribo:mRNA}, \qquad \text{Ribo. Binding,}$$

$$\text{Ribo:mRNA} + \text{AA} \underset{\text{aa}_{\text{Kd}}}{\overset{\text{aa}_{\text{F}}}{\rightleftharpoons}} \text{AA:Ribo:mRNA}, \qquad \text{AA Binding,}$$

$$\text{AA:Ribo:mRNA} + \text{AGTP} \underset{\text{TL}_{\text{n,Kd}}}{\overset{\text{TL}_{\text{n,F}}}{\rightleftharpoons}} \text{AA:AGTP:Ribo:mRNA}, \qquad \text{ATP-GTP Binding,}$$

$$\text{AA:AGTP:Ribo:mRNA} \xrightarrow{k_{\text{TL}}} \text{Ribo:mRNA}_{\text{term}} + \text{protein} + \text{AGMP}, \qquad \text{TL Elongation,}$$

$$\text{AA:AGTP:Ribo:mRNA} \xrightarrow{(Lp-1)k_{\text{TL}}} \text{Ribo:mRNA} + \text{AGMP}, \qquad \text{Consumption,}$$

$$\text{Ribo:mRNA}_{\text{term}} \xrightarrow{\text{ribo}_{\text{term}}} \text{Ribo} + \text{mRNA}, \qquad \text{Ribo. Release.}$$

*mRNA degradation reactions*

$$\text{RNAse} + \text{mRNA} \underset{\text{RNAse}_{\text{Kd}}}{\overset{\text{RNAse}_{\text{F}}}{\rightleftharpoons}} \text{RNAse:mRNA}, \qquad \text{RNAse Binding,}$$

$$\text{RNAse:mRNA} \xrightarrow{\text{RNAse}_{\text{cat}}} \text{RNAse}, \qquad \text{mRNA Degradation,}$$

$$\text{RNAse} + \text{Ribo:mRNA} \underset{\text{RNAse}_{\text{Kd}}}{\overset{\text{RNAse}_{\text{F}}}{\rightleftharpoons}} \text{RNAse:Ribo:mRNA}, \qquad \text{RNAse Binding,}$$

$$\text{RNAse:Ribo:mRNA} \xrightarrow{\text{RNAse}_{\text{cat}}} \text{RNAse} + \text{Ribo}, \qquad \text{mRNA Degradation,}$$

$$\text{RNAse} + \text{AA:Ribo:mRNA} \underset{\text{RNAse}_{\text{Kd}}}{\overset{\text{RNAse}_{\text{F}}}{\rightleftharpoons}} \text{RNAse:AA:Ribo:mRNA}, \qquad \text{RNAse Binding,}$$

$$\text{RNAse:AA:Ribo:mRNA} \xrightarrow{\text{RNAse}_{\text{cat}}} \text{RNAse} + \text{AA} + \text{Ribo}, \qquad \text{mRNA Degradation,}$$

$$\text{RNAse} + \text{AA:AGTP:Ribo:mRNA} \underset{\text{RNAse}_{\text{Kd}}}{\overset{\text{RNAse}_{\text{F}}}{\rightleftharpoons}} \text{RNAse:AA:AGTP:Ribo:mRNA}, \qquad \text{RNAse Binding,}$$

$$\text{RNAse:AA:AGTP:Ribo:mRNA} \xrightarrow{\text{RNAse}_{\text{cat}}} \text{RNAse} + \text{AGTP} + \text{AA} + \text{Ribo}, \qquad \text{mRNA Degradation.}$$

*Miscellaneous reactions*

In addition to the reactions above we consider two other mechanisms. A deGFP maturation reaction, and a simple mechanism to account for the cessation of TX-TL's ability to create more mRNA and protein. In the latter mechanism, the energy species AGTP is modeled to constantly degrade and regenerate up to a time $\tau_{\text{atp}}$, after which a Simbiology event sets the regeneration rate to zero, leading to rapid first order degradation of the remaining AGTP. The reversible reaction is shown as the second equation in Equations S2.1, and the trajectory of the AGTP degradation is shown in the bottom right panel of Figure 2.

$$
\begin{aligned}
&\mathrm{GFP} \xrightarrow{k_{\mathrm{mat}}} \mathrm{GFP}^*, && \text{GFP maturation,} \\
&\mathrm{AGMP} \underset{\delta_{\mathrm{atp}}}{\overset{\alpha_{\mathrm{atp}}}{\rightleftharpoons}} \mathrm{AGTP}, && \text{ATP-GTP regeneration-degeneration.}
\end{aligned}
\tag{S2.1}
$$

#### S2.3 Parameter Inference

There are a total of 26 parameters in this model, many of which are non-identifiable [7]. Both this large dimensionality and non-identifiability make sampling from the parameter distribution computationally difficult, possibly due to the multi-modal nature of the parameter densities [3]. We found that reducing the dimensionality by fixing some of the parameters allowed for the MCMC algorithm to eventually converge towards the desired distribution for the remaining parameters. We tried fixing different combinations of parameters, and explored the ability of the parameter inference algorithm to fit the model to the data. The various combinations of parameters we tried fixing are described in Table I. We found that using a 48-physical core Amazon EC2 instance (Intel) or a 16-physical core AMD Ryzen 9 processor, along with different combinations of MCMC mixing ("temperature") and step size hyperparameter values, we were able to fit the model to the data when the number of free parameters was at most 15 to 18.

Fixing the values of some parameters required identifying a point in the parameter space where the models fit the data at least approximately. Thus, we needed a 'nominal' parameter point that lay inside the set of (output-indistinguishable [9]) points at which the model fit the data, so that we could fix some of the parameter values, and infer the distributions of the remaining parameters (conditioned on the fixed values). This had to be done without solving the entire high-dimensional parameter inference problem itself (since that is what we were trying to find an estimate of in the first place).

To find such a point, we began by reducing the dimension of the search space from 26 to 23 by manually fixing three parameters: the deGFP maturation, AGTP degradation and AGTP regeneration rates. The values chosen were such that the dynamics of the toolbox were insensitive to the exact values, while at the same time gave GFP maturation and AGTP degradation-regeneration dynamics that were in approximate agreement with dynamics observed in the literature [6]. The maturation rate only affected the relationship between the unfolded and folded deGFP trajectories, and a workable value was found without difficulty. The AGTP regeneration rate was nonzero before time $\tau_{\mathrm{atp}}$, where it served to replenish AGTP, and its exact value did not matter as long as it did this quickly enough. The degradation rate set the time it took for the AGTP to disappear once the regeneration stopped, and could be directly compared to the profile in [6]. Furthermore, it had a direct effect on the duration the mRNA (resp. protein) trajectories were nonzero (resp. increasing), and therefore was also informed by the constitutive expression data. With these three parameter fixed, we attempted to infer the distribution of the resulting 23 parameter vector using the data from [2] (without the 'composite data' modifications described in Supplementary Section S2.1 above).

This smaller 23-dimensional inference problem was still computationally intractable, and the MCMC algorithm had to be terminated before it converged to the stationary distribution. Inspection of the individual trajectories generated by some of the sampled points led to the discovery of a candidate with trajectories qualitatively similar to the experimental data. After some tuning of the parameter values, we settled on a parameter point at which the model behavior and data looked similar, and it turned out that using this point as the nominal point worked for the purposes of the iterative parameter inference procedure, which we describe next.

As described in the main text, once the nominal parameter point had been picked, we tried different combinations of parameters to estimate (with the remaining fixed to the nominal values). Table I shows a summary of these test runs, and Supplementary Figures S2-S4 show the corresponding parameter distributions, and the experimental data and model fit trajectories.

In Run 1, only the forward reaction rates of the reversible reactions were fixed, and we verified that most of the values of the nominal point lay approximately inside the ranges of values where the respective nominal distributions had significant probability mass. This run, however, had too many (nineteen) free parameters, leading to extremely slow convergence. We terminated the run early, and moved to Run 2, which had fewer free parameters, and was able to fit the models to the data more rapidly.

More generally, we fixed the forward rate parameters (which set the timescale of reversible reactions) or the parameters that were highly non-identifiable [7, 1]. Examples of non-identifiable parameters include the parameters associated with the binding of nucleotides to the DNA-polymerase complex, or the binding of amino acids to the mRNA-ribosome complex. After fixing these parameters, we found that the parameters that remained could be inferred efficiently.

The parameter inference performed in Runs 1 - 5 was on the data from Reference [2]. To infer parameters for the composite data (described in Section S2.1 above), we chose the parameter fixing profile in Run 3. The results of the fitting are shown in Figure 3 and Supplementary Figure S5.

### S3 Inferring Incoherent Feed-forward Loop Part Parameters

#### S3.1 Experimental Data

As described in the main text, we performed five part characterization experiments, and five IFFL perturbation experiments. These experiments are shown in Table II. The TX-TL reactions were performed according to established protocols, as described in the Materials and Methods section (Section VI) of the main text, and the plasmids were designed by screening variants in TX-TL, also described in that section. Supplementary Figure S6 below shows the part screening procedure described in Section VI.

#### S3.2 Model Equations

**Constitutive pLac expression model**

The reactions for the constitutive pLac promoter model are identical to the constitutive expression model in Section S2.2. The code to generate the model is almost exactly the same as well, and is shown below.

```
tube1 = txtl_extract('E1');
tube2 = txtl_buffer('E1');
tube3 = txtl_newtube('pLacdeGFP');
```

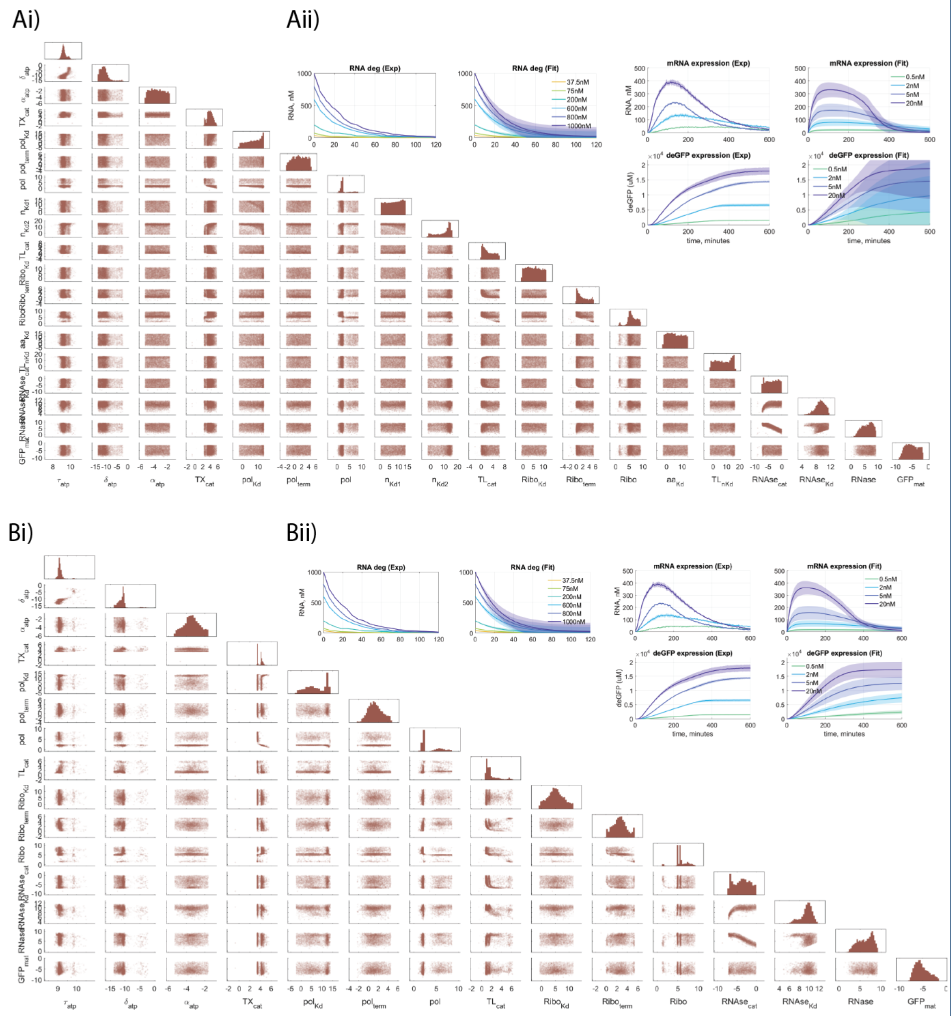

**Figure S2** Fitting trajectories and pairwise marginal posterior distributions for Runs 1 and 2 described in Table I.

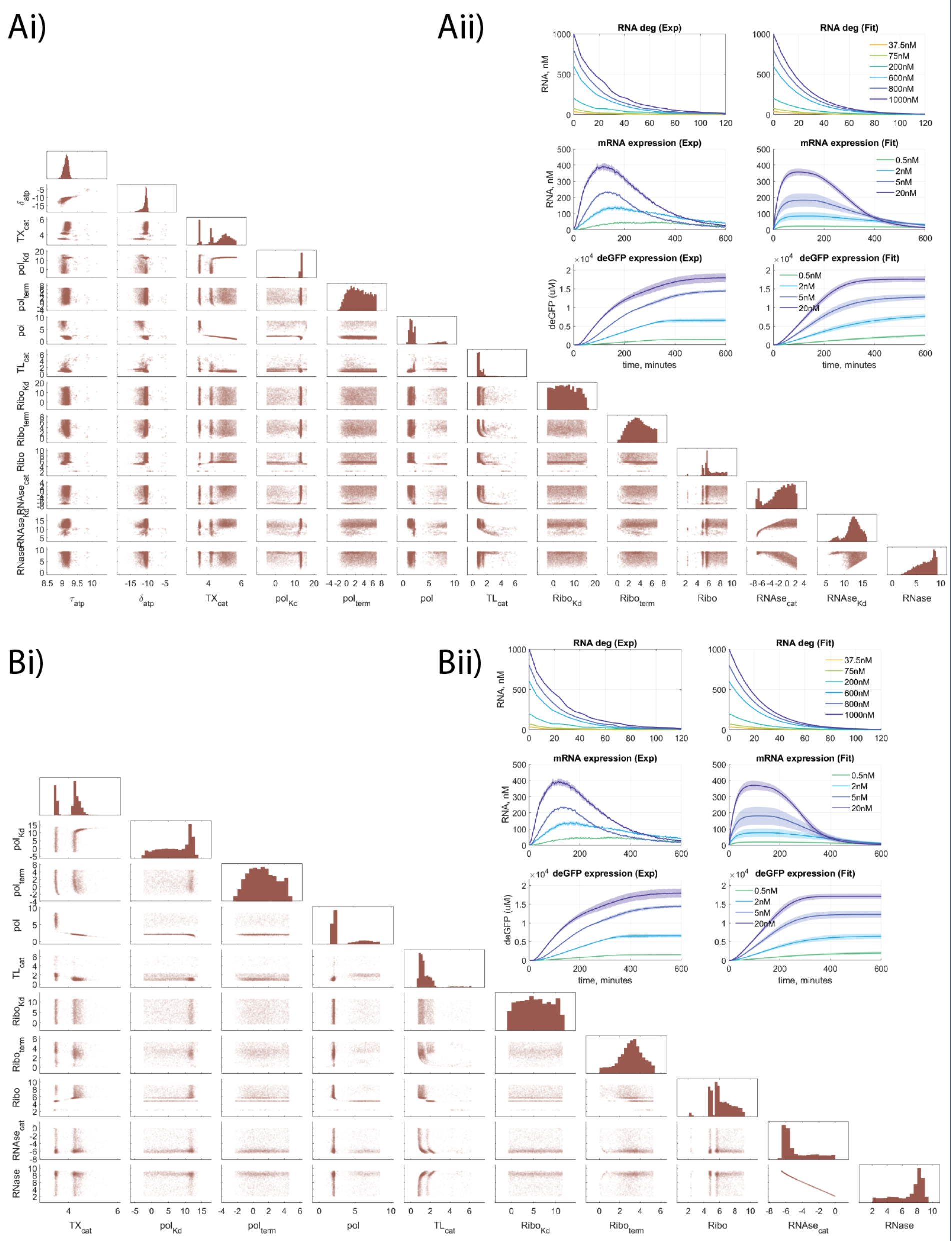


**Figure S3** Fitting trajectories and pairwise marginal posterior distributions for Runs 3 and 4 described in Table I.

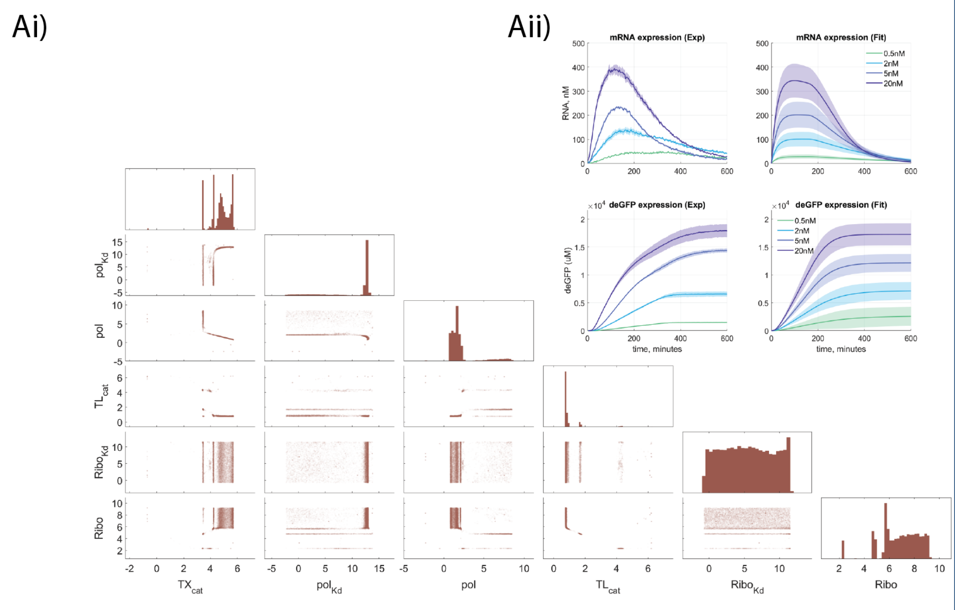


**Figure S4** Fitting trajectories and pairwise marginal posterior distributions for Run 5 described in Table I.

```
txtl_add_dna(tube3, 'plac(50)', 'utr1(20)', 'deGFP(1000)', 0, 'plasmid');
Model_obj = txtl_combine([tube1, tube2, tube3]);
simData = txtl_runsim(Model_obj, 8*60*60); % Simulate
txtl_plot(simData, Model_obj); % Plot
```

##### TetR repression system model

The code to generate the tet repression system is as follows.

```
tube1 = txtl_extract('E1');
tube2 = txtl_buffer('E1');
tube3 = txtl_newtube('pTetdeGFP_pLacTetR_aTc');
txtl_add_dna(tube3, 'ptet(50)', 'utr1(20)', 'deGFP(1000)', 0, 'plasmid');
txtl_add_dna(tube3, 'plac(50)', 'utr1(20)', 'tetR(1000)', 0, 'plasmid');
Model_obj = txtl_combine([tube1, tube2, tube3]);
txtl_addspecies(m, 'aTc', 0);
simData = txtl_runsim(Model_obj, 8*60*60); % Simulate
txtl_plot(simData, Model_obj); % Plot
```

The transcription, translation, mRNA degradation reactions are repeated for both the pTet-UTR1-deGFP and pLac-UTR1-TetR transcriptional units. The only reactions in addition to these (and the miscellaneous deGFP maturation and AGTP degradation-regeneration reactions) are the TetR dimerization and sequestration (induction) reactions,

$$\begin{aligned} 2\,\mathrm{TetR} &\underset{\mathrm{dim_{Kd}}}{\overset{\mathrm{dim_F}}{\rightleftharpoons}} \mathrm{TetR_{dimer}}, \\ \mathrm{DNA} + \mathrm{TetR_{dimer}} &\underset{\mathrm{rep_{Kd}}}{\overset{\mathrm{rep_F}}{\rightleftharpoons}} \mathrm{DNA{:}TetR_{dimer}}, \\ 2\,\mathrm{aTc} + \mathrm{TetR_{dimer}} &\underset{\mathrm{aTc_{Kd}}}{\overset{\mathrm{aTc_F}}{\rightleftharpoons}} \mathrm{aTc_2{:}TetR_{dimer}}. \end{aligned} \tag{S3.1}$$

##### LasR induction system model

The code to generate the LasR induction model is,

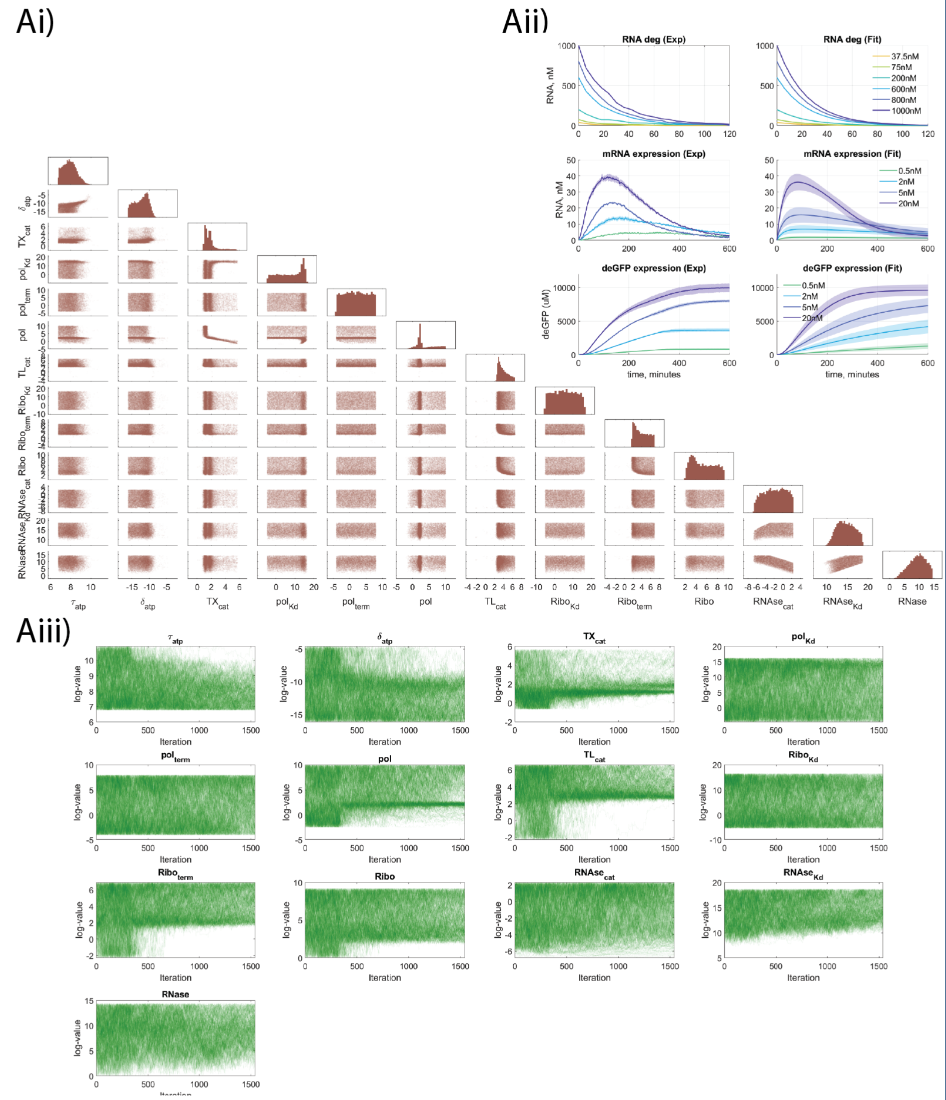

**Figure S5** Fitting trajectories, Markov chain trace plots and pairwise marginal posterior distributions for Run 3 performed on the composite data. The changes in the Markov chain plots at about 400 iterations are due to a change in the MCMC mixing parameter value that was employed as part of a simulated annealing procedure.

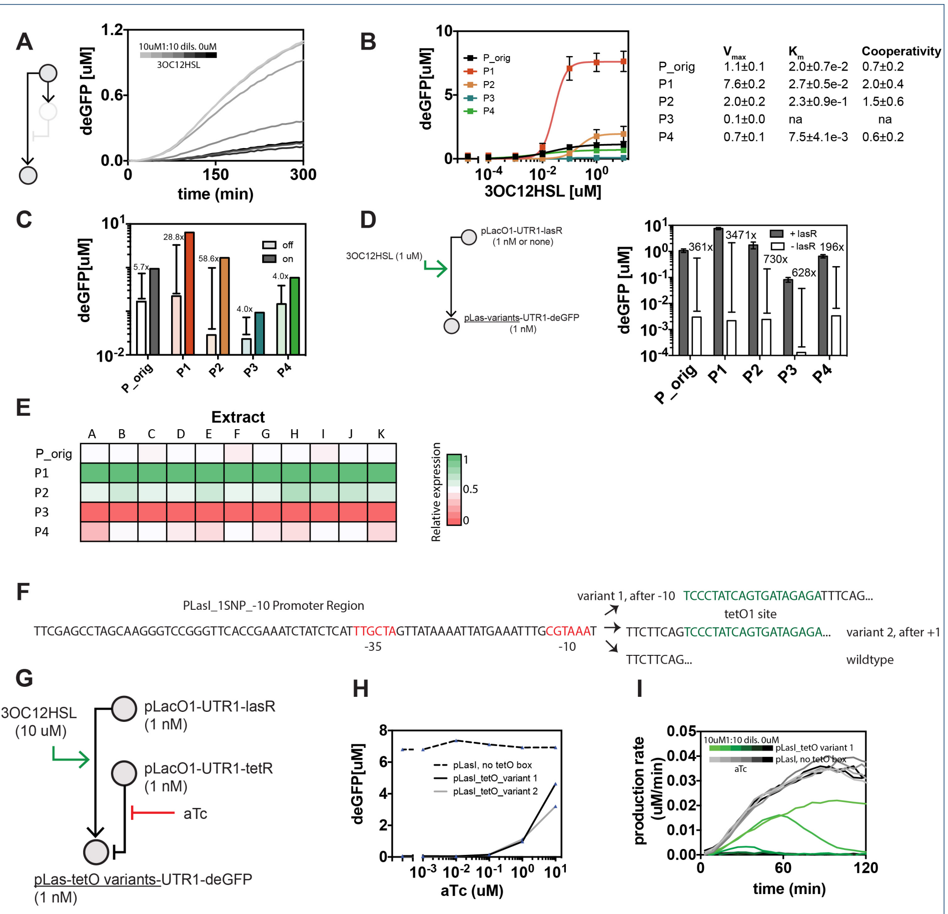


**Figure S6** Engineering a LasR activatable promoter and a las-tet combinatorial promoter. A) An LasR-activated promoter used in literature, pRsaL, is tested in TX-TL by varying 3OC12HSL. Circuit composed of 1 nM pLac-LasR plasmid and 1 nM pRsaL- deGFP (P_orig) plasmid. The mean of 3 independent experiments is shown. B) Alternative LasR-activated promoters are tested and characterized in TX-TL. Plotted is endpoint fluorescence at t = 300 minutes of 3 independent experiments at varying 3OC12HSL. Circuit composed of 1 nM pLac-LasR plasmid and 1 nM varying pLas promoter driving deGFP plasmid. C) Dynamic range of alternative promoters compared to pRsaL in off (no 3OC12HSL) vs. on (10uM 3OC12HSL), using data from B. D) Background expression of promoters in the absence of LasR in TX-TL. Each las promoter variant expressing deGFP is tested in TX-TL in the presence or absence of LasR. Plotted is endpoint expression after 5 hours subtracted for negative, with fold change difference. E) Comparison of promoter Vmax values at t = 180 minutes across different extracts. Each column is taken relative to the strongest and weakest expressing promoter. Extract A is used in this study. Extract A and D are an ExpressIQ strain, B-C and E-I are a BL21 Rosetta2 strain, J is a MGZ1 strain, and K a JS006 strain. F-I) De novo design of TetR-repressible pLas. F) Design of two placements of tetO operator sites, variant 1 and variant 2, compared to wild type. The difference is in the location of the tetO box. G) Circuit diagram for testing repression, where aTc is varied in the presence of a TetR-producing construct. H) Each promoter variant along with wild type is tested for repression ability with varying aTc. Shown are endpoint fluorescence at t = 300 min. I) First derivative indicating production rate of promoters.

```
tube1 = txtl_extract('E1');
tube2 = txtl_buffer('E1');
tube3 = txtl_newtube('pLacLasR_pLasdeGFP');
txtl_add_dna(tube3, 'plac(50)', 'utr1(20)', 'lasR(1000)', 0, 'plasmid');
txtl_add_dna(tube3, 'plas(50)', 'utr1(20)', 'deGFP(1000)', 0, 'plasmid');
Model_obj = txtl_combine([tube1, tube2, tube3]);
txtl_addspecies(m, 'OC12HSL', 0);
simData = txtl_runsim(Model_obj, 8*60*60); % Simulate
txtl_plot(simData, Model_obj); % Plot
```

Once again, the transcription, translation and mRNA degradation for each of the two transcriptional units (pLac-UTR1-LasR and pLas-UTR1-deGFP), along with the AGTP degradation-regeneration and deGFP maturation reactions, are as described for constitutive expression. The only modified mechanics in the model involve the binding of the activator to the promoter, and the resulting transcription via the activated DNA. This is described in the reactions below.

*Activation reactions:*

$$\text{RNAP} + \text{DNA} \underset{\text{pol}_{\text{Kd}}}{\overset{\text{pol}_{\text{F}}}{\rightleftharpoons}} \text{RNAP:DNA}, \qquad \text{Leaky Pol. Binding,}$$

$$\text{LasR\_protein} + \text{3OC12HSL} \underset{\text{3OC12}_{\text{Kd}}}{\overset{\text{3OC12}_{\text{F}}}{\rightleftharpoons}} \text{LasR\_protein:3OC12HSL}, \qquad \text{Inducer Binding,}$$

$$\text{LasR\_protein:3OC12HSL} + \text{DNA} \underset{\text{pLas}_{\text{TF, Kd}}}{\overset{\text{pLas}_{\text{TF, F}}}{\rightleftharpoons}} \text{DNA:LasR\_protein:3OC12HSL}, \qquad \text{Activation,}$$

$$\text{DNA:LasR\_protein:3OC12HSL} + \text{RNAP} \underset{\text{pLas-pol}_{\text{TF, Kd}}}{\overset{\text{pLas-pol}_{\text{TF, F}}}{\rightleftharpoons}} \text{RNAP:DNA:LasR\_protein:3OC12HSL}, \qquad \text{Pol. Binding Activated Prom.,}$$

$$\text{RNAP:DNA} + \text{LasR\_protein:3OC12HSL} \underset{\text{pLas}_{\text{TF, Kd}}}{\overset{\text{pLas}_{\text{TF, F}}}{\rightleftharpoons}} \text{RNAP:DNA:LasR\_protein:3OC12HSL}, \qquad \text{Activation,}$$

*Transcription reactions:*

$$\text{RNAP:DNA:LasR\_protein:3OC12HSL} + \text{AGTP} \underset{\text{n}_{\text{Kd1}}}{\overset{\text{n}_{\text{F1}}}{\rightleftharpoons}} \text{AGTP:RNAP:DNA:LasR\_protein:3OC12HSL},$$

$$\text{RNAP:DNA:LasR\_protein:3OC12HSL} + \text{CUTP} \underset{\text{n}_{\text{Kd1}}}{\overset{\text{n}_{\text{F1}}}{\rightleftharpoons}} \text{CUTP:RNAP:DNA:LasR\_protein:3OC12HSL},$$

$$\text{CUTP:RNAP:DNA:LasR\_protein:3OC12HSL} + \text{AGTP} \underset{\text{n}_{\text{Kd2}}}{\overset{\text{n}_{\text{F2}}}{\rightleftharpoons}} \text{CUTP:AGTP:RNAP:DNA:LasR\_protein:3OC12HSL},$$

$$\text{AGTP:RNAP:DNA:LasR\_protein:3OC12HSL} + \text{CUTP} \underset{\text{n}_{\text{Kd2}}}{\overset{\text{n}_{\text{F2}}}{\rightleftharpoons}} \text{CUTP:AGTP:RNAP:DNA:LasR\_protein:3OC12HSL},$$

$$\text{CUTP:AGTP:RNAP:DNA:LasR\_protein:3OC12HSL} \xrightarrow{k_{\text{TX}}} \text{RNAP:DNA}_{\text{term}} + \text{mRNA}, \qquad \text{TX Elongation,}$$

$$\text{CUTP:AGTP:RNAP:DNA:LasR\_protein:3OC12HSL} \xrightarrow{\left(\frac{Lm}{4}-1\right)k_{\text{TX}}} \text{RNAP:DNA}, \qquad \text{Consumption,}$$

$$\text{RNAP:DNA}_{\text{term}}\text{:LasR\_protein:3OC12HSL} \xrightarrow{\text{pol}_{\text{term}}} \text{RNAP} + \text{DNA} + \text{LasR\_protein:3OC12HSL}, \qquad \text{Pol. Release.}$$

#### S3.3 Parameter Inference

We adopted a sequential approach to estimating the part specific parameters for the IFFL. The overall trajectory of our exploration of the parameter space is summarized in Table III and Section IV

#### S3.4 Parameter Consensus Pattern

Next, we describe the parameter consensus pattern. I.e., we describe the mapping from the set of all parameters (both estimated and fixed) to the models they were involved in. Table S4 summarizes this consensus pattern. In this table, there are four unique models (described in Sections S2.2 andS3.2),

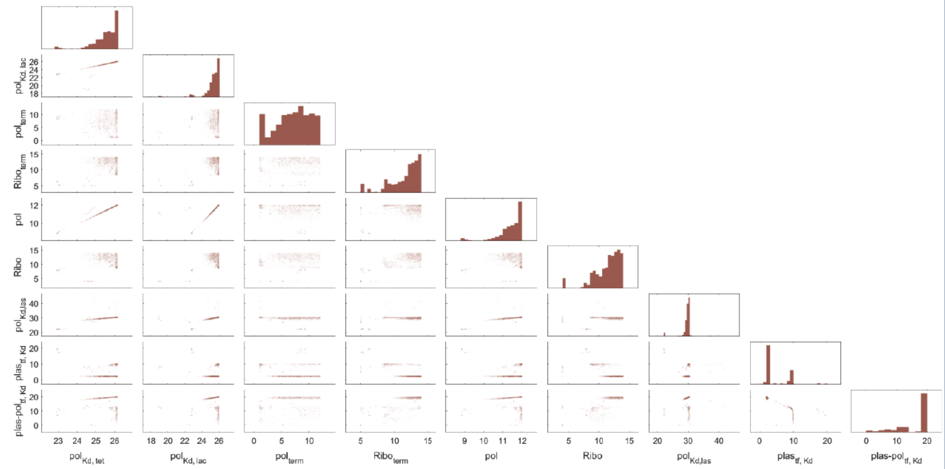

**Figure S7** Posterior distribution of the parameters for the IFFL part characterization Stage 2f (Table III). The corresponding experimental data and model fit trajectories are in Figure 4.

- vnprl: the generic constitutive expression model used in Stage 1,
- mlac: the lac constitutive expression model that is identical to the generic model, but with the lac promoter,
- mtet: the tet system model, which incorporates TetR mediated repression, and aTc mediated induction, and
- mlas: the las system model, which incorporates LasR and 3OC12 mediated activation.

Dosing and measured species describe the initial conditions and the outputs of the models respectively. The stages refer to those defined in Table III.

**Author details**

[1]Genome Institute of Singapore, 60 Biopolis Street, 138672, Singapore. [2]Computation and Neural Systems, California Institute of Technology, 1200 E. California Blvd, 91125 Pasadena, CA, USA. [3]Department of Bioengineering, Imperial College London, South Kensington, SW7 2AZ, London, UK. [4]Department of Bioengineering, Caltech. [5]Tierra Biosciences, West Berkeley, 94710, Berkeley, CA, USA. [6]Control and Dynamical Systems, Caltech.

**References**

1. O.-T. Chis, J. R. Banga, and E. Balsa-Canto, "Structural Identifiability of Systems Biology Models: A Critical Comparison of Methods," *PLoS ONE*, vol. 6, no. 11, p. e27755, Nov. 2011. [Online]. Available: http://dx.doi.org/10.1371/journal.pone.0027755
2. Dan Siegal-Gaskins, Z. A. Tuza, J. Kim, V. Noireaux, and R. M. Murray, "Gene Circuit Performance Characterization and Resource Usage in a Cell-Free "Breadboard"," *ACS Synthetic Biology*, vol. 3, no. 6, pp. 416–425, Jun. 2014. [Online]. Available: http://dx.doi.org/10.1021/sb400203p
3. D. Foreman-Mackey, D. W. Hogg, D. Lang, and J. Goodman, "emcee: The MCMC Hammer," *Publications of the Astronomical Society of the Pacific*, vol. 125, no. 925, pp. 306–312, Mar. 2013, arXiv: 1202.3665. [Online]. Available: http://arxiv.org/abs/1202.3665
4. C. Y. Hu, J. D. Varner, and J. B. Lucks, "Generating Effective Models and Parameters for RNA Genetic Circuits," *ACS Synthetic Biology*, Jun. 2015. [Online]. Available: http://dx.doi.org/10.1021/acssynbio.5b00077
5. E. Karzbrun, J. Shin, R. H. Bar-Ziv, and V. Noireaux, "Coarse-Grained Dynamics of Protein Synthesis in a Cell-Free System," *Physical Review Letters*, vol. 106, no. 4, p. 048104, Jan. 2011. [Online]. Available: http://link.aps.org/doi/10.1103/PhysRevLett.106.048104
6. V. Noireaux, R. Bar-Ziv, and A. Libchaber, "Principles of cell-free genetic circuit assembly," *Proceedings of the National Academy of Sciences*, vol. 100, no. 22, pp. 12 672–12 677, 2003. [Online]. Available: http://www.pnas.org/content/100/22/12672.short
7. A. Raue, C. Kreutz, T. Maiwald, J. Bachmann, M. Schilling, U. Klingmüller, and J. Timmer, "Structural and practical identifiability analysis of partially observed dynamical models by exploiting the profile likelihood," *Bioinformatics*, vol. 25, no. 15, pp. 1923–1929, Aug. 2009. [Online]. Available: https://academic.oup.com/bioinformatics/article/25/15/1923/213246/Structural-and-practical-identifiability-analysis

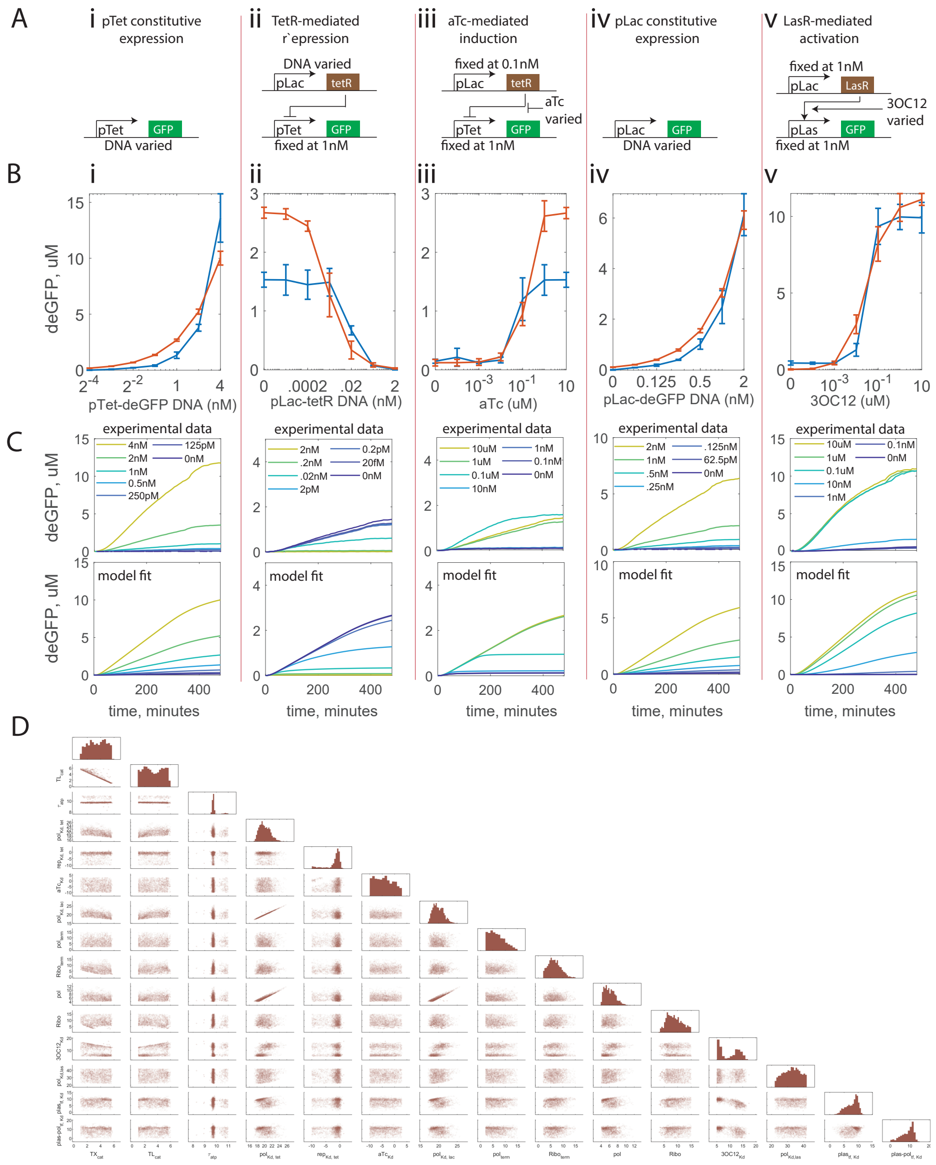


**Figure S8** Part Characterization and Parameter Fitting for the Incoherent Feed-Forward Loop, similar to Figure 4, but using the parameter fixing profile of Stage 2d in Table III. (A) Schematics describing the five part characterization experiments. (B) Endpoint curves (mean, standard error at 480 min) corresponding to the experimental data (blue, n = 3) and corresponding parameter fitting trajectories (orange, n = 50, sampled from the posterior parameter distribution and simulated). The posterior distributions were generated by fitting the full time-course trajectories to the data (C). (D) Pairwise marginal posterior parameter distributions resulting from the MCMC procedure.

**Table S4** Consensus parameter inference strategy.

| Model | vnprl | mtet | mtet | mtet | mLac | mlas |
|---|---|---|---|---|---|---|
| Dosing (initial cond.) | deGFP DNA | deGFP DNA | deGFP DNA TetR DNA | deGFP DNA TetR DNA aTc | deGFP DNA | deGFP DNA LasR DNA 3OC12HSL |
| Measured species (outputs) | deGFP mRNA | GFP | GFP | GFP | GFP | GFP |
| $n_{F1}$ | ✓ | ✓ | ✓ | ✓ | ✓ | ✓ |
| $n_{Kd1}$ | ✓ | ✓ | ✓ | ✓ | ✓ | ✓ |
| $n_{F2}$ | ✓ | ✓ | ✓ | ✓ | ✓ | ✓ |
| $n_{Kd2}$ | ✓ | ✓ | ✓ | ✓ | ✓ | ✓ |
| $ribo_F$ | ✓ | ✓ | ✓ | ✓ | ✓ | ✓ |
| $aa_F$ | ✓ | ✓ | ✓ | ✓ | ✓ | ✓ |
| $aa_{Kd}$ | ✓ | ✓ | ✓ | ✓ | ✓ | ✓ |
| $TL_{n,F}$ | ✓ | ✓ | ✓ | ✓ | ✓ | ✓ |
| $TL_{n,Kd}$ | ✓ | ✓ | ✓ | ✓ | ✓ | ✓ |
| $RNAse_F$ | ✓ | ✓ | ✓ | ✓ | ✓ | ✓ |
| $k_{mat}$ | ✓ | ✓ | ✓ | ✓ | ✓ | ✓ |
| $\alpha_{atp}$ | ✓ | ✓ | ✓ | ✓ | ✓ | ✓ |
| $k_{TX}$ | ✓ | ✓ | ✓ | ✓ | ✓ | ✓ |
| $pol_{term}$ | ✓ | ✓ | ✓ | ✓ | ✓ | ✓ |
| $pol_{init}$ | ✓ | ✓ | ✓ | ✓ | ✓ | ✓ |
| $k_{TL}$ | ✓ | ✓ | ✓ | ✓ | ✓ | ✓ |
| $ribo_{Kd}$ | ✓ | ✓ | ✓ | ✓ | ✓ | ✓ |
| $ribo_{term}$ | ✓ | ✓ | ✓ | ✓ | ✓ | ✓ |
| $ribo_{init}$ | ✓ | ✓ | ✓ | ✓ | ✓ | ✓ |
| $RNAse_{Kd}$ | ✓ | ✓ | ✓ | ✓ | ✓ | ✓ |
| $RNAse_{cat}$ | ✓ | ✓ | ✓ | ✓ | ✓ | ✓ |
| $RNAse_{init}$ | ✓ | ✓ | ✓ | ✓ | ✓ | ✓ |
| $\delta_{atp}$ | ✓ | ✓ | ✓ | ✓ | ✓ | ✓ |
| $\tau_{atp}$ | ✓ | ✓ | ✓ | ✓ | ✓ | ✓ |
| $pol_F$ (p70) | ✓ | | | | | |
| $pol_{Kd}$ (p70) | ✓ | | | | | |
| $pol_F$ (pLac) | | | | | ✓ | |
| $pol_{Kd}$ (pLac) | | | | | ✓ | |
| $pol_{Kd}$ (pTet) | | ✓ | ✓ | ✓ | | |
| $pol_F$ (pTet) | | ✓ | ✓ | ✓ | | |
| $rep_{Kd}$ | | | ✓ | ✓ | | |
| $rep_F$ | | | ✓ | ✓ | | |
| $aTc_{Kd}$ | | | | ✓ | | |
| $aTc_F$ | | | | ✓ | | |
| $dim_{Kd}$ | | | ✓ | ✓ | | |
| $dim_F$ | | | ✓ | ✓ | | |
| $pLas_{TF,\ F}$ | | | | | | ✓ |
| $pLas_{TF,\ Kd}$ | | | | | | ✓ |
| $3OC12_{Kd}$ | | | | | | ✓ |
| $3OC12_F$ | | | | | | ✓ |
| $pLas\text{-}pol_{TF,\ F}$ | | | | | | ✓ |
| $pLas\text{-}pol_{TF,\ Kd}$ | | | | | | ✓ |
| $pol_{TF,\ F}$ | | | | | | ✓ |
| $pol_{TF,\ Kd}$ | | | | | | ✓ |


8. V. Singhal and R. M. Murray, "Transforming Data Across Environments Despite Structural Non-Identifiability," in *2019 American Control Conference (ACC)*, Jul. 2019, pp. 5639–5646.
9. E. Walter and Y. Lecourtier, "Global approaches to identifiability testing for linear and nonlinear state space models," *Mathematics and Computers in Simulation*, vol. 24, no. 6, pp. 472–482, Dec. 1982. [Online]. Available: http://www.sciencedirect.com/science/article/pii/0378475482906450
